# A Role for Exaptation in Sculpting Sexually Dimorphic Brains from Shared Neural Lineages

**DOI:** 10.1101/2025.06.04.657833

**Authors:** Aaron M. Allen, Megan C. Neville, Tetsuya Nojima, Faredin Alejevski, Stephen F. Goodwin

## Abstract

Sex differences in behaviours arise from variations in female and male nervous systems, yet the cellular and molecular bases of these differences remain poorly defined. Here, we take an unbiased, single-cell transcriptomic approach to uncover how sex shapes the adult *Drosophila melanogaster* brain. We show that sex differences do not result from large-scale transcriptional reprogramming but through fine-tuning of otherwise shared developmental templates via the sex-differentiating transcription factors Doublesex and Fruitless. We reveal, with unprecedented resolution, the extraordinary genetic diversity within these sexually dimorphic cell types and find birth order represents a novel axis of sexual differentiation. Neuronal identity in the adult reflects spatiotemporal patterning and sex-specific survival, with female-biased neurons arising early and male-biased neurons arising late. This pattern reframes dimorphic neurons as “paralogous” rather than “orthologous”, suggesting sex leverages distinct developmental windows to build behavioural circuits and highlights a role for exaptation in diversifying the brain.

## INTRODUCTION

The nervous systems of sexually reproducing species diverge between the sexes to produce distinct behavioural repertoires^1^. As neuronal diversification in animals relies on the intrinsic mechanisms of spatial and temporal patterning, each sex must differentially act upon these mechanisms to establish neural circuits that meet their own behavioural needs^2^. Understanding how chromosomal sex informs the relationships between transcriptomic signatures and specific anatomical, physiological, and functional properties of cells in the nervous system is critical to understanding sexual behaviour.

The vinegar fly, *Drosophila melanogaster*, offers an ideal model to explore these questions due to its compact yet highly specialized central nervous system, well-characterized sex determination pathway and rich repertoire of sexually dimorphic behaviours. In *Drosophila*, sex is determined early in embryogenesis via an X chromosome counting mechanism, which activates the master regulator *Sex-lethal* (*Sxl*) in female^3–5^. *Sxl* triggers a hierarchical alternative splicing cascade, via *transformer* (*tra*), that culminates in the expression of two key transcription factors, Doublesex (Dsx) and Fruitless (Fru), which initiate many irreversible cell-autonomous sexual differentiation events (reviewed in^6,7^).

The expression of Dsx and Fru in the nervous system begins post-embryogenesis, sculpting sex-specific neural circuits required for adult sexual behaviour^8–11^. Neurons in *Drosophila* arise from neural stem cells, called neuroblasts, that divide to generate paired hemilineages, which represent the fundamental developmental and anatomical units of the central nervous system^12–17^. The selective expression of Dsx and Fru within specific hemilineages governs sex-specific survival and connectivity of dimorphic neurons, laying the groundwork for innate sexual behaviours (reviewed in^6^). However, the precise transcriptional landscape underlying the developmental specification of individual dimorphic neurons remains unclear.

Existing single-cell RNA sequencing (scRNA-seq) atlases have provided valuable overviews of cellular diversity across *Drosophila* tissues^18^. The central brain is an elaborately complex tissue critical for integrating sensory information, internal state, and behavioural outputs. To date, no single study has achieved sufficient cellular depth to detect the breadth of dimorphisms between the sexes across the brain. To overcome this limitation, we generated a high-depth, sexed single-cell transcriptomic atlas of the adult *Drosophila* central brain. By integrating our newly generated sex-specific data with multiple publicly available scRNA-seq datasets, we constructed a comprehensive meta-central brain neuron atlas encompassing 329,466 neurons^19^. In our companion manuscript, we show that neuronal identity is primarily defined by hemilineage, and secondarily by birth order within hemilineage^19^. Here, we build on this framework to investigate how sex shapes neuronal identity across the central brain, uncovering its influence at an unprecedented level of resolution.

Our analysis reveals that sexually dimorphic neuronal populations overwhelmingly correlate with the expression of *dsx* and *fru*, reinforcing their roles as key determinants of neuronal sexual identity. We generated *dsx*+ and *fru*+ neuron sub-atlases, enabling us to resolve distinct transcriptomic cell types, which we linked to anatomically defined cell types using genetic intersectional tools, allowing an understanding of transcriptomic diversity in a spatial and functional context. Our data uncover a novel relationship between sex and developmental birth order within hemilineage: female-biased neurons are typically born earlier, while male-biased neurons are typically born later. This finding suggests that sexually dimorphic neurons within a hemilineage often arise through developmental shifts in the survival of cells distinguished by their transcriptional identities and birth dates, rather than through a one-to-one correspondence of type between female and male counterparts, with females simply having fewer cells of that type. This reconceptualizes traditional models of sexual differentiation and offers a new framework for studying the evolution of sexually dimorphic neural circuits.

## RESULTS

### Sex-specific transcriptomic diversity in the adult *Drosophila* central brain

To investigate how chromosomal sex shapes transcriptional diversity in the nervous system, we generated a sexed single-cell transcriptomic atlas of the *Drosophila melanogaster* adult central brain^19^. We integrated our data with multiple independent central brain-containing single-cell datasets, resulting in a comprehensive meta-central brain neuronal atlas with a cellular depth of coverage of 9.8x and comprising 329,466 neurons (Figure 1A; see^19^ for details).

**Figure 1.**
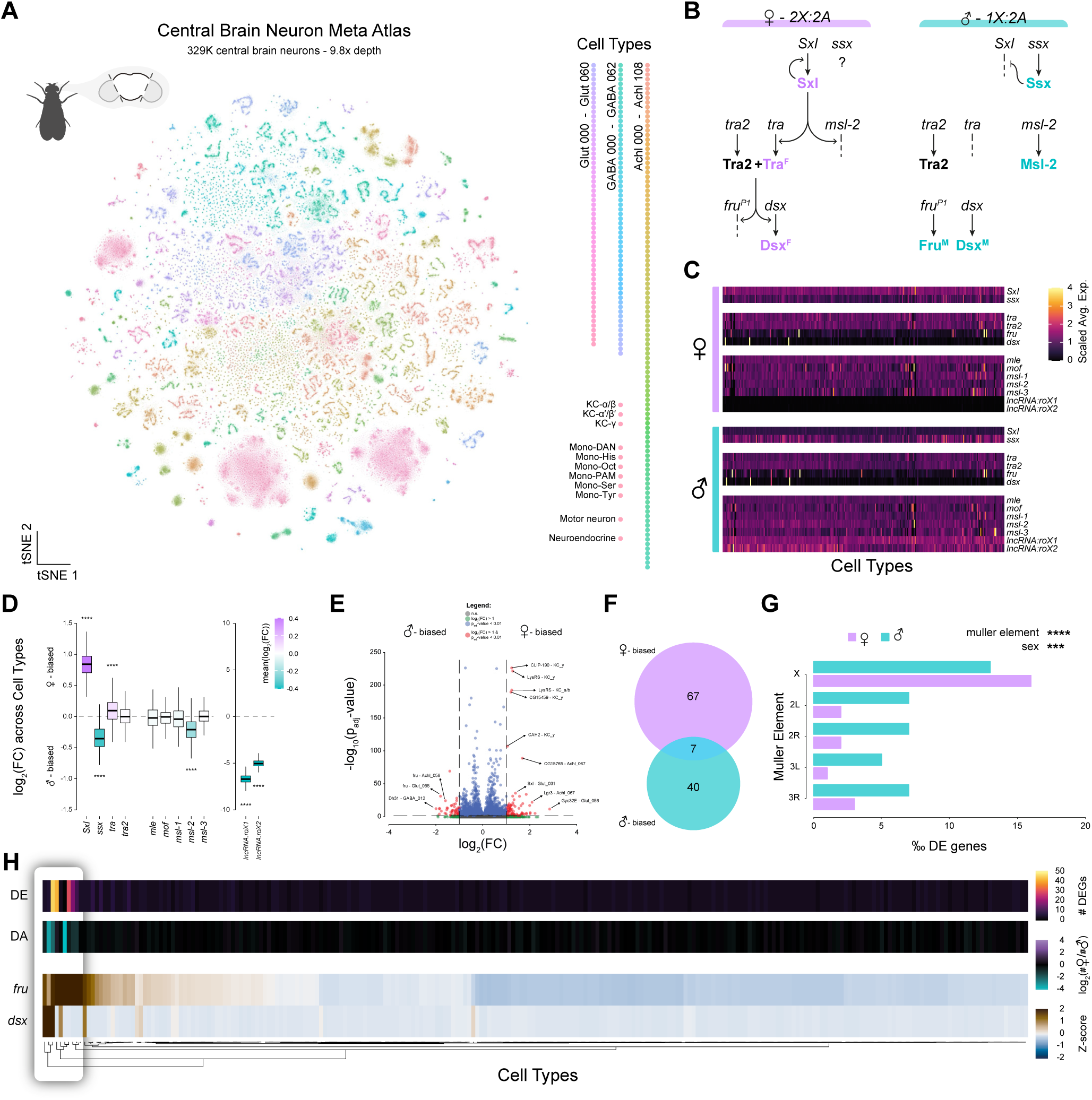
Sex dimorphism in *Drosophila* central brain at single-cell resolution. (**A**) t-distributed stochastic neighbour embedding (tSNE) of 329,466 *Drosophila* central brain neurons, representing 9.8-times depth coverage. Cell types are defined by unique colour spectrums. Fast-acting neurotransmitter types: acetylcholine (Achl), GABA, and glutamate (Glut); Monoaminergic types: dopaminergic (DAN), Histidine (His), octopaminergic (Oct), serotonergic (Ser), Tyraminergic (Tyr), and dopaminergic positive allosteric modulators (PAM); Kenyon cell (KC) types, motor neurons, and endocrine cells. (Data from companion paper^19^). (**B**) Schematic depiction of the sex determination dosage compensation hierarchy in *Drosophila melanogaster*. (**C**) Heatmap of scaled average expression of sex determination and dosage compensation genes across all neuronal cell types of the central brain in females (top) and males (bottom). (**D**) Boxplots showing log_2_ fold-change (FC) of sex-biased sex determination (left) and dosage compensation (right) gene expression across all cell types in the central brain. Bonferroni adjusted Chi-squared p-value less than 0.0001 (****). (**E**) Volcano plot of differentially expressed genes exhibiting male-biased (left) and female-biased (right) expression identified in all neuronal cell types across the central brain. Dashed lines indicate statistical significance thresholds; p_adj_-value<0.01, log_2_(FC)>1. (**F**) Euler diagram representing the number of differentially expressed genes amongst cell types unique to females, males, or shared between the sexes (albeit in different cell types). (p_adj_-value<0.01, log_2_(FC)>1). (**G**) The distribution of differentially expressed (DE) genes across Muller elements (chromosomal arms) between females and males. Bonferroni adjusted Chi-squared p-value less than 0.001 (***) and 0.0001 (****). (**H**) The number of differentially expressed genes (DE, top), differential abundance of cell numbers between the sexes (DA, middle), and z-scored expression of *dsx* and *fru* (bottom) across all cell types are shown. Differential expression was filtered as in E. Cell types are hierarchical clustered based on the expression of *dsx* and *fru*. Boxed region highlights cell types with the highest DE, DA, and *dsx* or *fru* expression.

To explore sex differences in transcriptionally distinct cell types within central brain neurons, we first examined the expression of genes involved in the sex determination hierarchy (SDH) and dosage compensation (Figure 1B). The SDH governs somatic sexual differentiation, and most of its members were ubiquitously expressed across all neuronal cell types in both sexes (Figure 1C), including the master regulator *Sxl*, its translational repressor *sister-of-Sex-lethal*^20–22^ (*ssx*), and the splicing factors *tra* and *transformer-2* (*tra2*), whose widespread expression has recently been confirmed throughout the body^23^. In contrast, the transcription factors *dsx* and *fru* exhibited cell-type-restricted expression, marking the first level of spatially specific transcriptional regulation underlying sexual differentiation in the central brain (Figure 1C). Analysis of dosage compensation genes (Figure 1B) revealed that the male-specific long non-coding RNAs *lncRNA:roX1* and *lncRNA:roX2* were ubiquitously expressed in males, while other dosage compensation genes were ubiquitously detected in both sexes (Figure 1C). *roX1* and *roX2* were the only genes exhibiting both ubiquitous and sex-specific expression. Several SDH and dosage compensation pathway components showed sex-biased expression across all neuronal cell types: *Sxl* and *tra* displayed significant female-biased expression, while *ssx* and *msl-2* were significantly male-biased (Figure 1D). Notably, three genes – *Sxl*, *tra*, and *msl-2 –* were more highly expressed in the sex that produces the corresponding functional protein product (Figure 1B, 1D).

### Differential gene expression between the sexes in central brain neurons

We next examined sex-biased gene expression across the central brain in each independent neuronal cell type (Figure 1E). Applying stringent statistical criteria (see STAR Methods), we identified 114 differentially expressed genes (DEGs): 74 female-biased and 47 male-biased, 7 of which were female-biased in one cell type and male-biased in another (Figure 1E–F). Most of these genes (87%) were differentially expressed in only a single cell type, although the specific cell type varied for each gene. This low number of sex-biased genes identified in the central brain (∼1% of expressed genes), along with the modest magnitude of their differential expression, is striking, especially when compared to other tissues^18^. Gene ontology enrichment analysis revealed significant overrepresentation of terms related to transmembrane signalling, cyclic nucleotide-mediated pathways, and behaviours associated with mating and courtship (Figure S1A), suggesting that sex differences may influence circuit-level properties such as neuronal excitability, synaptic plasticity, and hormonal responsiveness. Our data support a model where sex subtly influences the transcriptome of mature neurons, suggesting functional dimorphism arises from precise, cell-type-specific tuning within a shared framework, not from dramatic transcriptional reprogramming across the brain. This, however, does not rule out the possibility of larger-scale transcriptional divergences during development.

Analysis of the genomic distribution of DEGs revealed a marked enrichment of sex-biased transcripts on the X chromosome relative to individual autosomal arms, with this bias being especially pronounced among female-biased genes (Figure 1G). This pattern, consistent with prior observations^18,24–26^, supports the notion that the X chromosome serves as a favourable genomic context for the emergence and regulation of sex-biased gene expression^24–26^. In male *Drosophila*, global dosage compensation mechanisms elevate X-linked gene expression to approximate that of the two X chromosomes in females. Interestingly, we identified *Myc*, a broadly female-biased transcription factor residing on the X chromosome, which appears to escape dosage compensation in males and has been implicated in the activation of *Sxl* in females (Figure S1B-D)^5,27,28^.

Sex differences in gene expression do not need to be large to exert significant biological effects, and small but widespread sex-biased expression patterns may accumulate to generate significant dimorphic traits^29^. When we lowered our thresholds to capture all DEGs with any fold-change, we observed numerous ubiquitously expressed genes showing subtle yet consistent sex-biased expression across multiple neuronal populations (Figure S1B-C). We validated the robustness of this bias across all datasets within our meta-atlas, excluding genes whose sex-biased expression was confined to individual datasets (Figure S1D; Figure S2). The broad expression of many SDH genes throughout the central brain (Figure 1C) indicates that genes such as *Sxl* and *tra* may influence widespread, low-level patterns of sex-biased expression independently of *dsx* and *fru*, as has been reported in other tissues^23,30,31^. Intriguingly, among the ubiquitously sex-biased genes, a pronounced trend emerged with the majority of ribosomal (129/164) and mitochondrial (81/89) genes exhibiting a trend of male-biased expression, suggesting potential sex-specific metabolic demands in neurons (Figure S1E). However, these trends approach the detection limits of our approach and may be influenced by technical factors.

Finally, we investigated the distribution of sex differences across neuronal populations of the central brain. To do this, we considered two measures – the number of differentially expressed genes per cell type (using stringent filtering) and the differential abundance (the number of female cells compared to the number of male cells) of each cell type (Figure 1H). Most cell types showed little to no difference in these metrics, with only a few exhibiting dramatic sex differences in the number of genes differentially expressed and differential abundance. When exploring what distinguishes these cell types, we found they showed strong enrichment for *dsx* or *fru*. These findings suggest that the strongest sex differences in the brain are highly regional, with *dsx* and *fru* serving as robust markers of neuronal populations where sex differences are most pronounced, consistent with their well-established roles in sexual differentiation (reviewed in^6^). Accordingly, we refer to *dsx+* and *fru+* neurons as the transcriptionally sexually dimorphic contingent of the central brain and to neurons lacking expression of both genes as transcriptionally sexually monomorphic, while acknowledging that low-level and widespread differences are present throughout the brain.

### Distinct transcriptional signatures define *doublesex*+ neuronal subtypes

We next explored the transcriptional diversity of *dsx*-expressing neurons. We extracted and subclustered all 2,108 *dsx*+ cells from our meta-central brain neuron atlas (Figure 2A). The resulting *dsx+* atlas, representing 9.8-fold depth of coverage, revealed six broad transcriptionally distinct neuronal subpopulations (Figure 2B) in both sexes (Figure 2C). To correlate these subpopulations with anatomically defined *dsx*+ cell types in the central brain, we leveraged established cell number representation and neurotransmitter usage to predict their identities (Figure 2D-E). Further molecular characterization revealed subtype-specific expression of transcription factors and transmembrane receptors, indicating distinct regulatory and physiological profiles that may underlie sex-specific circuit functions (Figure 2F).

**Figure 2.**
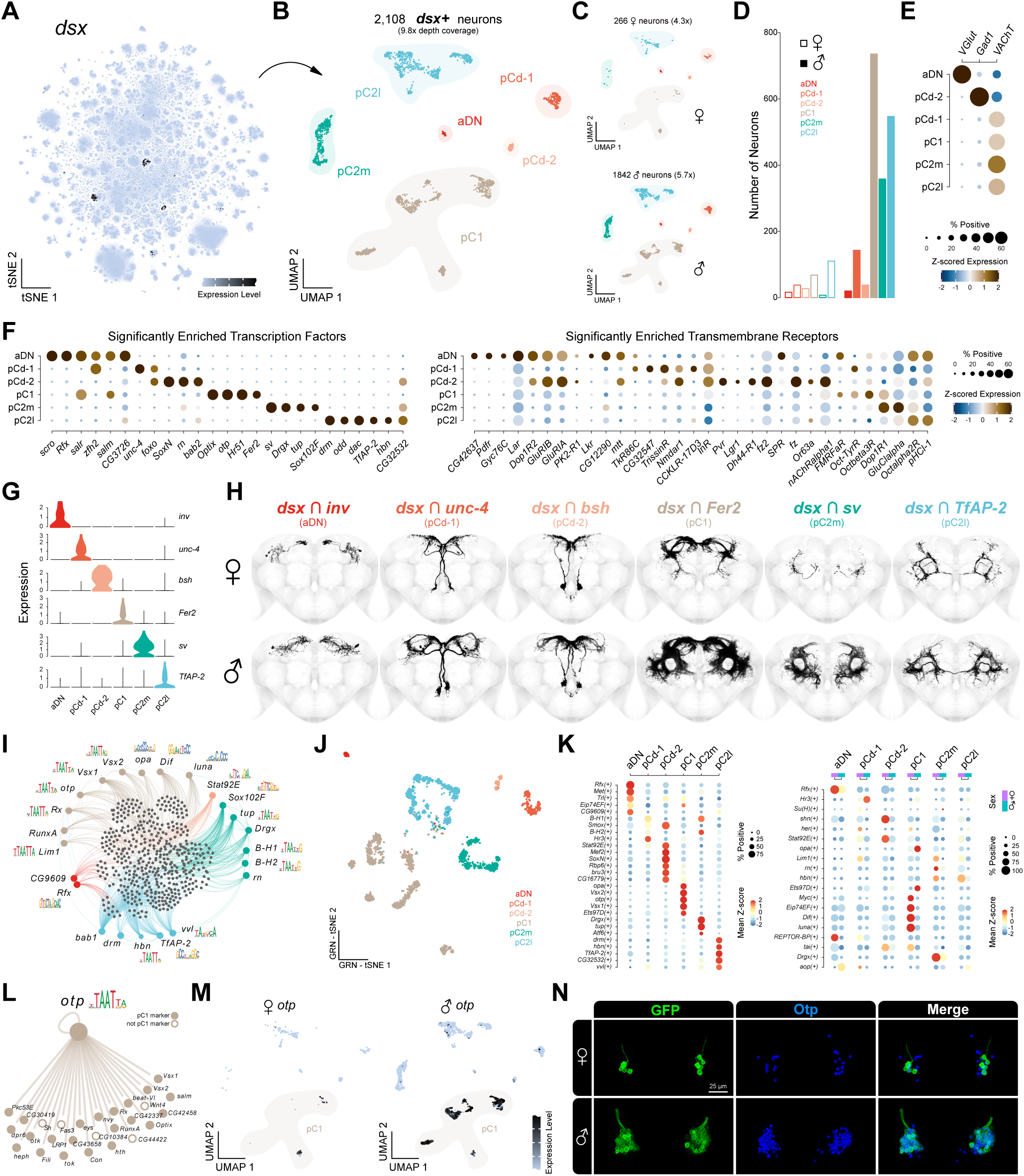
Transcriptional diversity of *doublesex* neurons in the central brain. (**A**) tSNE of *dsx* expression across central brain neurons. (**B**) Uniform manifold approximation and projections (UMAP) of subclustered *dsx+* neurons, with transcriptionally defined subtypes annotated. (**C**) UMAP of *dsx+* neurons in females (top) and males (bottom), representing 4.3x and 5.7x depth of coverage, respectively. (**D**) Bar chart of the number of *dsx+* neurons across transcriptionally defined central brain subtypes between females and males. (**E**) Dot plot showing expression of key biomarkers for acetylcholine (*VAChT*), glutamate (*VGlut*), and GABA (*Gad1*) across *dsx+* neuronal subtypes. (**F**) Dot plot of the expression of significantly enriched transcription factors (left) and transmembrane receptors (right) across *dsx+* neuronal subtypes. (**G**) Violin plot showing expression of key transcription factors labelling distinct *dsx+* neuronal subtypes. (**H**) Whole-mount immunofluorescence showing neuronal populations in the central brain identified via genetic intersection between *dsx* and key transcription factors (*inv, unc-4, bsh, Fer2, sv,* and *TfAP-2*). Images have been segmented, full expression patterns in Figure S3C. (**I**) Network diagram illustrating putative regulatory relationships between GRNs identified based on SCENIC analysis across *dsx+* neurons. Nodes are coloured based on GRN expression across *dsx*+ subtypes. (**J**) GRN-tSNE of *dsx+* neuronal subtypes based SCENIC GRN analysis, coloured by transcriptionally defined cell types. (**K**) Dot plots of significantly enriched regulons across *dsx+* neuronal subtypes (left), between the sexes (right). (**L**) *otp* GRN identified in *dsx*+ pC1 neurons. Gene nodes are coloured based on their identity as pC1 cluster markers. (**M**) UMAPs showing the expression of *otp* in females (left) and males (right) *dsx+* neurons. pC1 subtype cells are highlighted. (**N**) Immunofluorescence showing *dsx+* pC1 neurons (GFP, left), Otp expression (blue, middle) and co-expression (merged, right). pC1 soma and hemilineage-associated axonal tracks segmented from *dsx^Gal^*^4^>mCD8::GFP.

To validate these predicted identities, we employed a split-Gal4 strategy to genetically intersect *dsx* expression with subtype-specific transcription factors (Figure 2G-H; Figure S3A–C). This approach successfully labelled all six *dsx+* cell types and yielded cell counts consistent with prior anatomical estimates (Figure S3D)^10,32^. Thus, every cell of a given type is represented in these intersections, unifying their transcriptional identity with anatomical identity and providing a powerful and unprecedented toolkit for future functional studies investigating *dsx*+ neuronal populations.

Next, we used single-cell regulatory network inference and clustering (SCENIC) to identify gene regulatory networks (GRNs) that specify each *dsx+* subtype (Figure 2I–M). GRN-based subclustering closely paralleled transcriptional-based clustering (Figure 2J), underscoring the robustness of regulon architecture in defining neuronal identity. Many cell type-enriched regulons corresponded to *dsx*+ subtype-specific transcription factors (Figure 2F), such as *otp* and *Vsx1/2* in pC1, *Drgx* and *tup* in pC2m, and *TfAP-2* and *drm* in pC2l (Figure 2K). We further examined the *otp* regulon, which was selectively enriched in pC1 neurons. Genes within this regulon were among the most specific pC1 markers (Figure 2L), implicating Otp as a core determinant of this key behavioural hub’s identity, connectivity, and function. Immunohistochemical analysis confirmed Otp expression in female and male pC1 neurons (Figure 2M, N), consistent with its predicted regulatory role. These findings suggest that sex-specific circuit assembly is guided by Dsx and lineage-specific transcription factors such as Otp that shape neuronal subtype identity and connectivity.

### *fruitless*+ neurons developmental diverge between the sexes

To investigate the role of *fru* in sex differences in central brain neurons, we generated a *fru+* central brain neuron atlas by sub-clustering all *fru*-expressing cells (Figure 3A-B). Kenyon cells and monoaminergic neurons were analyzed separately (Figure S4). Using stringent expression thresholds, we identified 13,295 *fru*+ neurons. The cellular representation of *fru* in our atlas falls between previous estimates derived from antibody-based methods and *fru*-Gal4 reporter approaches^8,33^. In our companion paper^19^, we propose that the anatomical identity of transcriptionally defined cell types in our atlas approximates developmental hemilineage. Based on this, we estimate that *fru* is expressed within at least 57% of central brain hemilineages, consistent with its role as a temporal transcription factor marking subpopulations that reflect birth order^19^.

**Figure 3.**
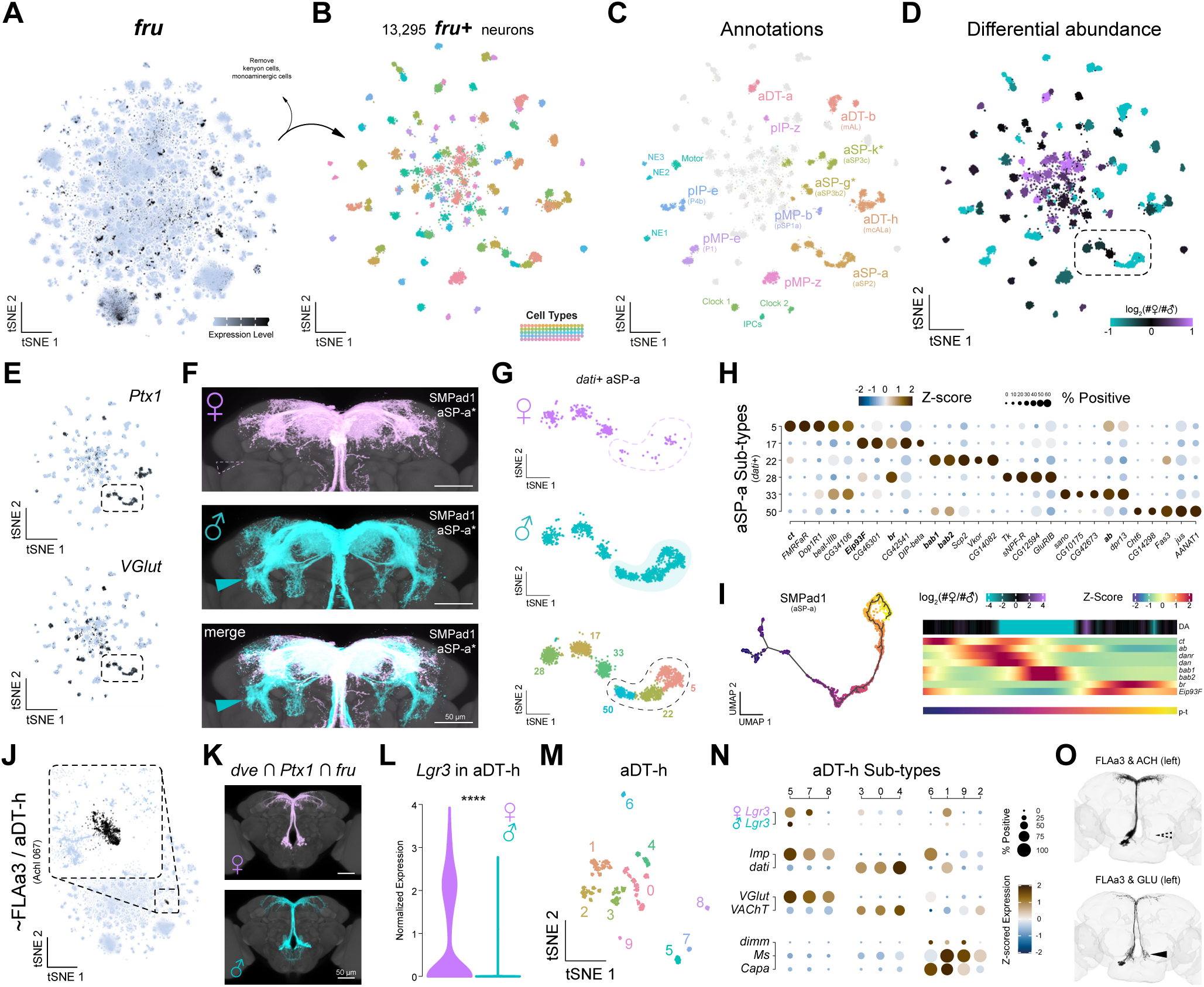
*fru*+ meta-analysis reveals sexually diverse cell types in the adult central brain. (**A**) tSNE of *fru+* expression across the *Drosophila* central brain atlas. (**B**) tSNE of 13,295 *fru+* neurons subclustered and coloured by transcriptionally unique cell types. Kenyon cells and monoaminergic cells were removed for clarity (see Figure S4). (**C**) Annotated tSNE of anatomically defined *fru*+ neuronal cell types. (**D**) tSNE of differential abundance in cell numbers between the sexes in *fru*+ cell types, coloured by log-transformed female-to-male cell number ratio. (**E**) tSNEs of *Ptx1* and *VGluT* expression in *fru*+ neurons, with overlapping expression highlighted by dashed boxes. (**F**) Immunofluorescence images of *Ptx1 ∩ VGlut* neurons in female (magenta, top) and male (blue, middle) central brains, with merged overlay (white, bottom). Solid blue arrowhead indicates male-specific neurites, while dashed magenta arrowhead marks their absence in females. Expression patterns were segmented; full patterns in Figure S7. (**G**) Zoomed-in region of the *dati*+ (late-born) aSP-a cell type within the *fru*+ central brain tSNE, showing female (top) and male (middle) cell distribution across transcriptionally distinct aSP-a subtypes (bottom). Male-specific subtypes (5, 22, and 50) are highlighted. (**H**) Dot plot of significantly enriched gene expression in *dati*+ aSP-a subtypes. (**I**) Pseudotime trajectory (left) of SMPad1 hemilineage (containing aSP-a neurons). Heatmap (right) of sex differences in cell abundance (top) and gene expression dynamics of transcriptional markers (middle) along pseudotime (p-t; bottom), showing key TFs defining early, middle, and late birth-order identities. (**J**) Central brain tSNE highlighting *fru*+ aDT-h/FLA3 hemilineage, corresponding to Achl 067 cell type (black), with zoomed-in inset (dashed box). (**K**) Immunofluorescence-based anatomical identification of aDT-h neurons using *dve*, *Ptx1*, and *fru* co-expression in female (top) and male (bottom) central brains. Expression patterns have been segmented, full patterns in Figure S7. (**L**) Violin plot of *Lgr3* expression in aDT-h neurons, showing significant female-bias (**** p < 0.0001). (**M**) UMAP sub-clustering of aDT-h neurons, coloured by transcriptionally distinct subtypes. (**N**) Dot plot of gene expression across aDT-h subtypes, highlighting subtype-specific expression of *Lgr3* (in females and males), early-vs. late-born markers (*Imp* and *dati*), neurotransmitter marker genes (*VGlut* and *VAChT*), and neurosecretory-specific TF *dimm* and neuropeptides (*Ms* and *Capa*). (**O**) EM reconstructions of FLAa3 neurons in one hemisphere of the central brain (FlyWire dataset), showing distinct cholinergic (ACH) and glutamatergic (Glut) populations, highlighting anatomically distinct neurites (arrowheads).

Using established markers, we annotated known neuronal populations, such as motor, neuroendocrine, and clock neurons (Figure 3C; Figures S5, S7). Other transcriptionally defined clusters were annotated through genetic intersectional strategies, linking them to anatomically characterized *fru*+ neurons (Figure S6; Figures S7). This approach provided a systematic pipeline for linking transcriptional profiles with the anatomical identities of *fru+* neurons. Notably, it identified both well-characterized populations (e.g., aSP-a and aDT-h) and previously unannotated *fru*+ cell types (pMP-z and pIP-z), highlighting its utility in uncovering the full diversity of sexually dimorphic neuronal populations.

Several *fru+* neuronal subtypes exhibited sex biases in cell numbers (Figure 3D). We focused on a prominent male-biased population corresponding to a central brain cell type with one of the highest numbers of differentially expressed genes (Figure 1H, cluster Glut 058). Using a double (*Ptx1* and *VGlut*) and triple (*fru*, *Ptx1*, and *VGlut*) genetic intersection strategy, we identified the glutamatergic *fru*+ aSP-a cell type (also known as aSP2; Figure 3E-G; Figure S7A,D,E), which has been previously described as one of the largest *fru*+ populations in the central brain^34^. Indeed, most subtypes of the SMPad1 hemilineage correspond to *fru*+ aSP-a neurons (Figure S7A). These neurons are localized in the superior medial protocerebrum (SMP) and form male-specific, characteristic ring-shaped arborizations in the lateral protocerebral complex (Figure 3F), a region densely innervated by *dsx*+ neurons (Figure 2H).

Within the aSP-a clusters, we identified six transcriptionally distinct subtypes, three of which are male-specific and likely contribute to the formation of male-specific arborizations (Figure 3E-H). Notably, these subtypes express *abrupt* and *beat-III* genes, which are known to harbour sex-biased enhancer elements that exhibit differential chromatin accessibility in aSP-a neurons^35^. Our companion paper^19^ shows that neuronal subtypes within a hemilineage are defined by temporal transcription factors associated with birth order that are retained in the adult. Building on this framework, we analyzed the entire SMPad1 hemilineage across pseudotime (Figure 3I). Male-specific subtypes emerged during a developmental window coinciding with the expression of *bab1* and *bab2*, temporal transcription factors that mark mid-to-late neurogenesis. This suggests that female neurons born during this window are eliminated via programmed cell death, a process that Fru is known to inhibit in males^36,37^.

Next, we focused on a cell type identified in the central brain as having dramatic sex-biased gene expression, which is entirely *fru*+ (Figure 1H, Figure 3J; Figure S7B). Genetic intersection of transcription factors co-expressed in this cell type, *Ptx1* and *dve*, identified the previously described *fru*+ aDT-h (also known as mcALa) neuronal cell type, corresponding to hemilineage FLAa3 (Figure 3K; Figure S7B and S7F)^38^. While this population is largely monomorphic^34^, we and others^39,40^ observed a slight male bias in cell number alongside distinct neurites in the gnathal ganglia that form male-specific, horsetail-like dendritic fields (Figure 3F). As expected, the *Insulin-like peptide 8* (*Ilp8*) receptor *Lgr3*, a Fru target previously shown to be repressed in aDT-h neurons^41^, exhibited strong female-biased expression (Figure 3L; Figure S7C).

Sub-clustering of aDT-h neurons revealed nine subtypes with distinct transcriptional signatures (Figure 3M). Unexpectedly, we observed female-specific *Lgr3* expression across three distinct subtypes: an early-born glutamatergic, a late-born cholinergic, and an endocrine subtype (Figure 3N). Exploring the FLAa3 hemilineage in the female connectome^42,43^ revealed two anatomically distinct subclasses corresponding to differential neurotransmitter usage, which reshapes our understanding of information flow within this hemilineage: a cholinergic subtype whose projections terminate in the dorsal-most part of the SMP, and a glutamatergic subtype that extends projections to the contralateral flange region, via the dorsal SMP, forming output sites in both neuropil regions (Figure 3O). Notably, the *Lgr3+* endocrine subtype co-expresses the neuropeptides *Myosuppressin* (*Ms*) and *Capability* (*Capa*) in both sexes (Figure 3N). These findings suggest that *Ilp8* signalling may modulate the release of multiple neuropeptides in a female-specific manner, influencing the physiological state of females^44,45^.

### Male-specific neuropeptide expression via Fruitless-dependent neuronal survival

While most central brain neuronal cell types displayed few or no DEGs between the sexes (Figure 1H), specific cell types exhibited dramatic sex differences in the expression of individual genes. A novel finding is the highly male-biased expression of the neuropeptide *Diuretic hormone 31* (*Dh31*) in single GABAergic neuronal population (GABA 012) (Figure 4A-C). Indeed, male-biased expression was consistently observed across all independent single-cell RNA-seq datasets within our central brain neuron atlas (Figure 4B), confirming its robustness and biological relevance. Dh31 is critical in regulating various physiological processes in *Drosophila* and is homologous to the vertebrate calcitonin gene-related peptide (CGRP)^46^.

**Figure 4.**
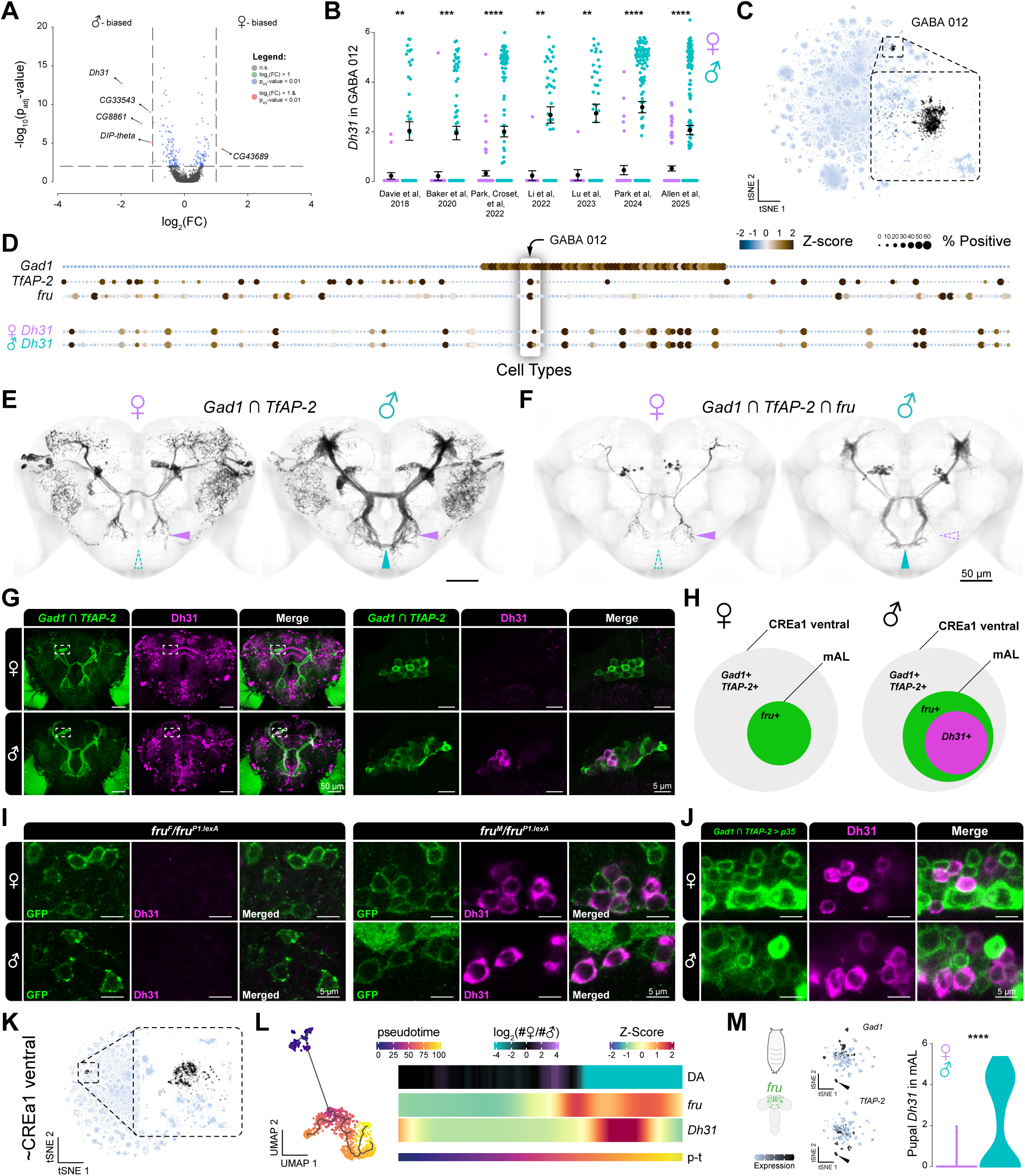
Male-specific expression of the neuropeptide Dh31 in the central brain. (**A**) Volcano plot of differentially expressed genes in females and males within the GABAergic 012. Dashed lines indicate statistical significance thresholds. (**B**) *Dh31* expression between females and males within the GABA 012 cluster, split by datasets. (** p < 0.01; *** p < 0.001; **** p < 0.0001, Bonferroni corrected Wilcoxon Signed-Rank Test). (**C**) tSNE highlighting GABA 012 (in black) within the neuronal central brain atlas, with zoomed- in inset (dashed box). (**D**) Dot plot of GABA 012 defining genes (*Gad1*, *TfAP-2*, *fru*; top) and *Dh31* (split by sex; bottom) across all neuronal clusters in the central brain. (**E**) Immunofluorescence of *TfAP-2* and *Gad1* intersected identifies the CREa1 ventral hemilineage. Arrowheads highlight anatomical differences in neurite projections between the sexes: magenta = female, blue = male, solid = present, dashed = absent. Expression patterns have been segmented, full patterns in Figure S8A. (**F**) Triple intersected *Gad1*, *TfAP-2*, and *fru* cells within the central brain identifies the sexually dimorphic mAL neuronal cluster. Expression patterns have been segmented, full patterns in Figure S8A. (**G**) Whole mount immunofluorescence images of *Gad1*∩*TfAP-2*>myr::GFP (green), anti-Dh31 (magenta), and merged (white). Female (top), male (bottom), whole brains (left), and dashed white box insets highlighting individual section soma (right). (**H**) Venn diagram of the CREa1 ventral hemilineage (grey), which encompasses all *fru*+ mAL cells in each sex (green), with a subset of male mAL neurons expressing Dh31 (magenta). (**I**) Dh31 expression in the mAL in hemizygous *fru^F^* females and males (*fru^F^*/*fru^P^*^1^*^.LexA^; left*), and in hemizygous *fru^M^* females and males (*fru^M^* /*fru^P^*^1^*^.LexA^; right*). Close-up of individual sections of mAL soma expressing myr::GFP (green), anti-Dh31 (magenta), and merged (white). Female (top) and male (bottom). Whole brains in Figure S8C. (**J**) Whole mount immunofluorescence of central brains with *Gad1*∩*TfAP-2* intersection expressing p35 and myr::GFP (green), anti-Dh31 (magenta), and merged (white). Female (top) and male (bottom). Whole brains are shown in Figure S8E. (**K**) Central brain neuron tSNE highlighting the mAL containing CREa1 ventral hemilineage (black), with zoomed-in inset (dashed box). (**L**) Pseudotime trajectory (left) of CREa1 ventral hemilineage. Heatmap (right) showing sex differences in cell abundance (top) and gene expression dynamics of *fru* and *Dh31* (middle) along pseudotime (p-t; bottom). (**M**) Schematic of mid-pupal, central brain, *fru* neurons (left), tSNEs (middle) highlighting *Gad1* (top) and *TfAP-2* (bottom) expression. Arrowhead highlights the mAL cell cluster. Violin plot of *Dh31* expression in each sex in the pupal mAL (right). (**** p < 0.0001, Wilcoxon Signed-Rank Test). Reprocessed data originally from Palmateer et al., 2023^52^ (see Figure S9).

To determine the anatomical identity of this GABAergic population, we identified the GABAergic marker *Gad1*, the transcription factors *TfAP-2*, and *fru* as robust markers of its transcriptional identity (Figure 4D). The genetic intersection of *TfAP-2* and *Gad1*, along with a triple intersection with *fru*, anatomically mapped this transcriptionally distinct cell type to the CREa1 ventral hemilineage (aka CREa1B), and its *fru+* subset mAL (Figure 4E-F, Figure S8A). mAL neurons represent a well-characterized sexually dimorphic population, known for their role in processing and integrating sex-pheromone cues in males, thereby ensuring that courtship behaviour is directed toward sexually receptive, species-appropriate mates^47–50^.

To validate the sex-specific expression of Dh31, we performed Dh31 immunostaining, which revealed male-specific expression in the CREa1 ventral neurons, specifically within the subset of *fru+* mAL neurons (Figure 4G-H; Figure S8B). To assess whether this sex difference is directly regulated by *fru*, we utilized *fru* splice mutants that induce constitutive splicing to either female (non-functional, *fru^F^*) or male (functional, *fru^M^*) isoforms^51^. Chromosomal females expressing the functional male isoform of *fru* (*fru^M^/fru^P^*^1^*^.lexA^*) exhibited Dh31 expression in mAL neurons, whereas chromosomal males expressing the non-functional female isoform of *fru* (*fru^F^/fru^P^*^1^*^.lexA^*) did not (Figure 4I; Figure S8C-D). These findings demonstrate that male-specific Dh31 expression depends on the functional male isoform of Fru (Fru^M^).

Fru protects male mAL neurons from programmed cell death (PCD), leading to fewer mAL neurons in females^36^. To investigate whether PCD influences *Dh31* expression in females, we inhibited cell death using the p35 apoptosis inhibitor, which restored Dh31 expression in female mAL neurons (Figure 4J; Figure S8E). This result indicates that the absence of Dh31 in female mAL neurons is due to female-specific PCD rather than male-specific *fru*-mediated transcriptional regulation of *Dh31*. Pseudotime analysis suggests that *fru* and *Dh31* co-positive, male-specific cells are born late in the CREa1 ventral hemilineage (Figure 4K-L). Furthermore, re-analysis of a pupal *fru+* scRNA-seq dataset^52^ (Figure S9) revealed that mAL neurons exhibit male-biased *Dh31* expression during mid-pupal development (Figure 4M). This suggests that female-specific cell death of *Dh31+* neurons occurs before 48 hours post-puparium formation. To explore potential downstream targets of local Dh31 release, we genetically intersected its receptor, *Dh31-R*, in both *dsx+* and *fru+* neurons, revealing broad expression in males. This suggests that Dh31 signalling coordinates sex-specific neuronal activity across multiple brain circuit elements (Figure S8F).

### Interplay between Dsx and Fru defines sexually dimorphic neuronal lineages

To investigate how *dsx* and *fru* collectively orchestrate the specification of sexually dimorphic neuronal identities, we merged their respective sub-atlases to construct a unified *dsx+* and *fru+* central brain atlas, encompassing all transcriptionally dimorphic neurons (Figure 5A–B). We then isolated cells within defined hemilineages – the developmental units comprising the central brain – that co-expressing *dsx* and *fru* to systematically explore their transcriptional relationships. Leveraging genetic intersectional tools, prior MARCM clonal analyses, and hemilineage-associated tract annotations from the connectome, we annotated and confirmed six hemilineages which co-express dsx and fru^32,34,37,38,40,42,43,53–55^.

**Figure 5.**
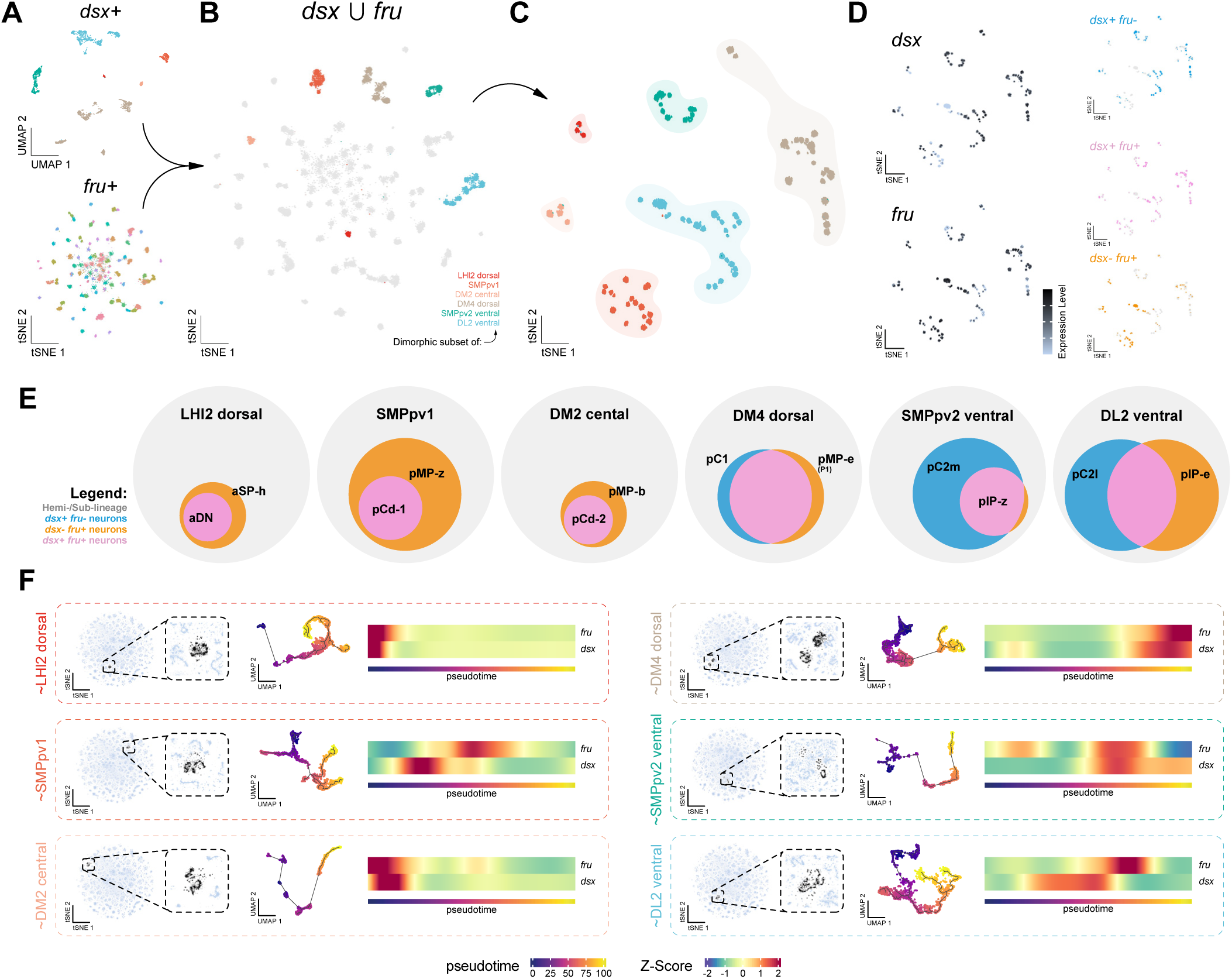
Developmental relationship between *dsx* and *fru* cell types in the central brain. (**A**) *dsx*+ (top) and *fru*+ (bottom) central brain neuronal sub-atlases. (**B**) tSNE of *dsx*+ and *fru*+ union central brain atlas (*dsx* U *fru*), coloured by populations with shared hemilineage identities (LHI2 dorsal, SMPpv1, DM2 central, DM4 dorsal, SMPpv2 ventral, DL2 ventral). (**C**) tSNE of subclustered *dsx* U *fru* neurons with shared hemilineage identities (indicated by colours). (**D**) tSNEs visualization of *dsx* and *fru* expression (left), with individual and co-expressed cells shown in different colours (right). (**E**) Venn diagrams depicting the relationship between *dsx*+ and *fru*+ expressing neurons within distinct hemilineages. (**F**) Pseudotime analyses of 6 *dsx*+, *fru*+ containing hemilineages in the central brain. tSNEs highlighted predicted hemilineages in reprocessed central brain atlas (left, see Star Methods). Pseudotime trajectories (middle) of hemilineages, and heatmaps (right) showing gene expression dynamics of *dsx* and *fru* along pseudotime.

Within each of these hemilineages, we resolved distinct neuronal subtypes co-expressing *dsx* and *fru*, in addition to subtypes expressing each transcription factor independently (Figure S11). This finding indicates that *dsx* and *fru* can act cooperatively and independently within hemilineages to diversify neuronal identities. To further investigate the inferred developmental dynamics between *dsx* and *fru*, we performed pseudotime analysis on extracted approximations of these six hemilineages (see STAR Methods), revealing striking lineage-specific variation in the onset and overlap of their expression (Figure 5F). The lack of chronological coordination of *dsx* and *fru* between the hemilineages suggests that the precise mechanisms by which Dsx and Fru act in concert to regulate neuronal identity and sex-specific differentiation during neurogenesis occur in a hemilineage-specific manner. Future experiments into these developmental dynamics will elucidate these patterns.

### Developmental timing and programmed cell death shape sex differences in pC1/P1 neurons

We observed considerable transcriptional heterogeneity among dimorphic neurons within each *dsx*+/*fru*+ hemilineage. To further explore this diversity, we focused on pC1/P1 cluster within the DM4 dorsal hemilineage, a critical integrative hub for social arousal in both females and males (Figure 6A). This cluster comprises anatomically and functionally distinct neuronal subtypes that differ in abundance and connectivity between the sexes, properties previously shown to depend on *dsx* and *fru*^10,56–65^. We identified nine broad transcriptionally distinct pC1/P1 subtypes (Figure 6B-C), and confirmed they correlated with morphologically distinct neuronal populations by genetic intersectional mapping (Figure 6D; Figure S12). This reinforces the finding that pC1/P1 subtypes are functionally specialised; indeed, a more refined analysis of this population reveals extensive transcriptional and cell type diversity, explored in detail in Figure S11.

**Figure 6.**
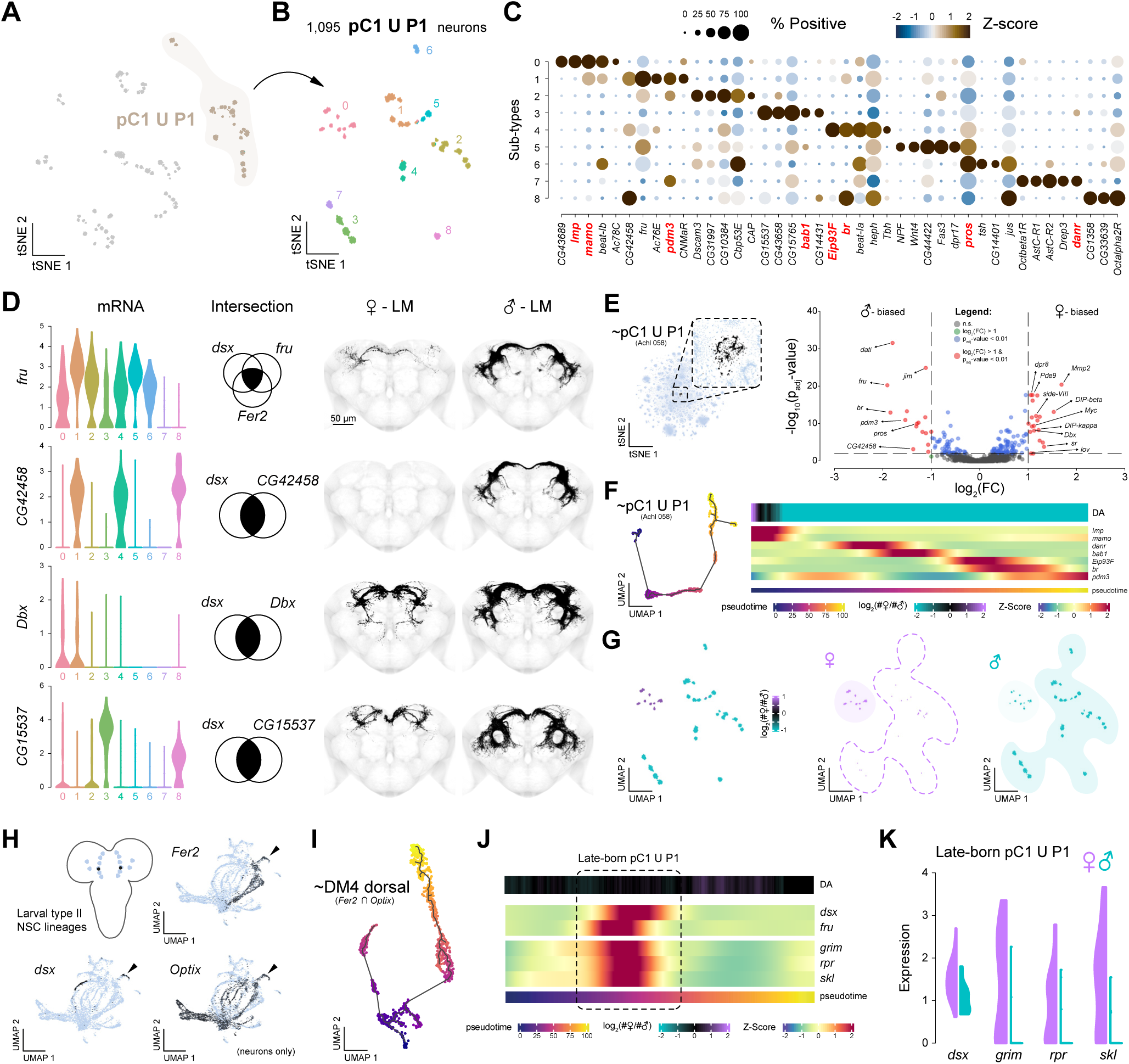
Genetically defining pC1/P1 neurons uncovers sex-specific cell death. (**A**) tSNE of *dsx* U *fru* neurons with shared hemilineage identities with the pC1/P1 subtype highlighted. (**B**) UMAP of 1,095 pC1/P1 neuron subclustering, coloured by transcriptionally distinct cluster identities. (**C**) Dot plot of top genes defining pC1/P1 subtypes. Developmental genes predicted to mark birth order within hemilineages^19^ are highlighted in red. (**D**) Violin plots of pC1/P1 subtype-defining gene expression (left). Genetic intersections identifying anatomically distinct cell types (middle). Immunofluorescence light microscopy (LM) images showing pC1/P1 neuronal populations identified via genetic intersection (right) in females and males. Images have been segmented; full expression patterns in Figure S12. (**E**) tSNE (left) highlighting the pC1/P1 containing cell type Achl 058 in the neuronal central brain atlas with zoomed-in inset (dashed box). Volcano plot (right) showing differentially expressed genes in Achl 058 cluster between sexes. Dashed lines indicate statistical significance thresholds. (**F**) Pseudotime trajectory (left) of Achl 058 cell type. Heatmaps (right) showing sex differences in cell abundance (top) and gene expression dynamics key genes defining early, middle and late birth-order identities (middle) along pseudotime (bottom). (**G**) UMAPs of differential abundance between the sexes in pC1 subtypes (left) and split by sex (right). Male-specific subtypes are highlighted within the dashed line. (**H**) Schematic of larval type II neuroblast lineages (top left). UMAPs of *Fer2*, *Optix*, and *dsx* expression in type II lineages (right, bottom) defining an approximation of the DM4 dorsal hemilineage. Reprocessed data originally from Michki et al., 2021^66^ and Rajan et al., 2023^67^. (**I**) Pseudotime analysis of DM4 dorsal hemilineage approximation (∼) defined by *Fer2* and *Optix* co-expression. (**J**) Heatmaps showing lack of sex differences in cell abundance (top) and gene expression dynamics of *dsx* and *fru* and cell death genes *grim*, *rpr*, and *skl* (middle) along pseudotime (bottom). (**K**) Split violin plot of *dsx* and cell death gene expression in post-embryonic larval pC1/P1 neurons in females and males.

When analyzing DEG between the sexes across the central brain, we found the cell type displaying the highest number of sex-biased genes as well as the most dramatic differential cell abundance between the sexes corresponds to pC1/P1 neurons (Achl 058; Figure 1H). Several of these differentially expressed genes encode temporal transcription factors that regulate neuron birth order and subtype identity during neurogenesis (Figure 6E). To explore the inferred developmental dynamics within the pC1/P1 neuronal population, we conducted a pseudotime analysis, revealing previously established temporal transcription factors^19^ expressed in distinct pseudotime windows (Figure 6F). Comparing these developmental trajectories with differential cell abundance between sexes, we found a female-biased temporal window which correlated with markers of early-born neurons (*Imp* and *mamo*), as well as a male-specific temporal window which correlated with mid- and late-born markers (Figure 6F). When examining the sex distribution among pC1/P1 subtypes, we found that all but one subtype was male-specific (Figure 6G). Thus, female and male pC1/P1 neurons arise from distinct temporal windows and represent non-overlapping neuronal subtypes.

To further resolve how developmental timing intersects with sexual identity, we analyzed larval scRNA-seq datasets from type II neuroblasts and their lineages, which includes the DM4 dorsal hemilineage that gives rise to pC1/P1 neurons^37,66,67^. We extracted DM4 dorsal cells and performed a pseudotime analysis, identifying a developmental window during which *dsx* and *fru* expression peaks (Figure 6H-J). While examining genes that co-varied with *dsx* and *fru* expression, we identified three key pro-apoptotic genes: *grim*, *rpr*, and *skl* (Figure 6J). Notably, the expression of these genes was female-specific in these post-embryonic, late-born pC1/P1 neurons (Figure 6K). Previous work demonstrated that both sexes experience PCD within the pC1/P1 cluster, with females experiencing more extensive neuronal loss than males, leading to highly sexually dimorphic numbers of neurons^37^ (Figure S3D). Our data refine this model, suggesting that female-specific PCD primarily eliminates late-born pC1/P1 neurons, whereas male-specific PCD targets early-born pC1/P1 neurons. Furthermore, we detected no sex difference in DM4 dorsal neuron abundance at late larval stages (Figure 6J), indicating that sex-biased PCD initiates during late larval development but completes post-puparium formation. Together, these data support a model in which birth order within a hemilineage dictates sexual fate via temporally gated, sex-specific PCD.

### Sexually dimorphic cells represent distinct sub-populations between the sexes

To examine the broader implications of birth order and sex-specific differentiation in the central brain, we first analysed differential cell abundance across *dsx*+ and *fru*+ neuronal populations (Figure 7A). We found female-biased neuronal types predominantly expressed the early temporal marker *Imp*, whereas male-biased types were enriched for the late-born marker *dati* (Figure 7B). Indeed, we observed a strong correlation between sex-biased cell numbers and these birth-order markers (Figure 7C), suggesting that temporal identity is broadly linked to sexual dimorphism across neuronal populations. Note that in our companion paper, we provide a detailed analysis of early- and late-born neurons in the central brain, revealing that these populations display distinct transcriptional signatures^19^.

**Figure 7.**
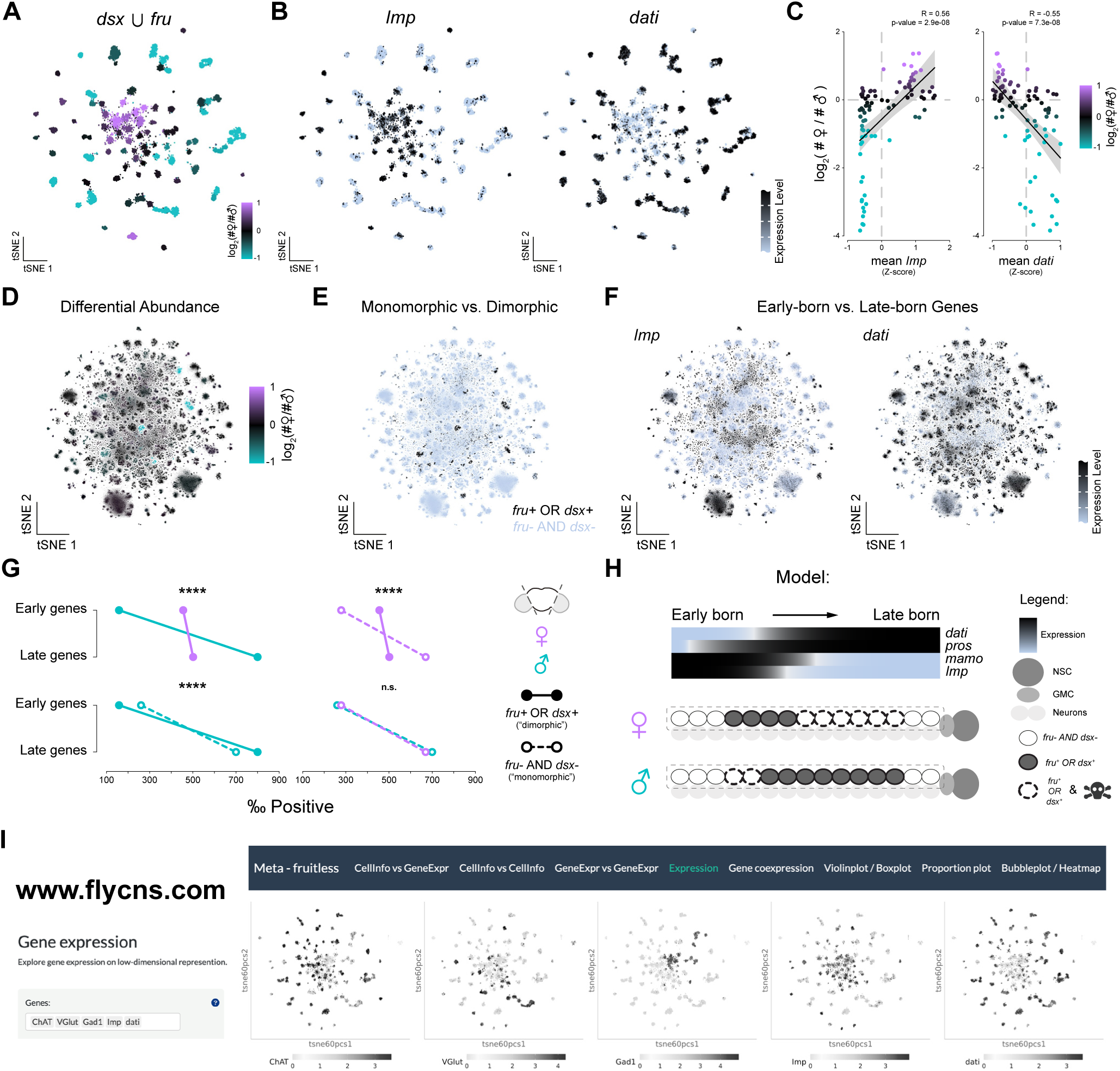
Sexually dimorphic neurons experience differential cell death. (**A**) tSNE of *dsx* U *fru* neurons showing differential abundance in cell numbers between the sexes across cell types. (**B**) tSNE of *Imp* (left) and *dati* (right) expression across *dsx* U *fru* neurons. (**C**) Scatter plots showing correlations of sex-biased cell numbers with *Imp* (left) and *dati* (right) expression across *dsx* U *fru* neuronal populations. (**D**) tSNE of differential abundance between the sexes across neuronal cell types in the central brain. (**E**) Central brain tSNE showing classification of neurons as either transcriptionally monomorphic or dimorphic based on *dsx* and *fru* expression. Neurons expressing *dsx* or *fru* (black) are classified as dimorphic, while neurons lacking both (blue) are monomorphic. (**F**) tSNEs of early-born (*Imp*, left) and late-born (*dati*, right) marker gene expression across the central brain neurons, indicating developmental timing differences in neuronal populations. (**G**) Per mille (‰) of cells expressing early genes (*Imp*, *mamo*) and late genes (*dati*, *pros*), for dimorphic cells (*fru+* OR *dsx+*, solid lines) and monomorphic cells (*fru-* AND *dsx-*, dashed lines) in females and males (see Star Methods). (**** p < 0.0001, n.s. = not significant, Bonferroni corrected Chi-squared test). (**H**) Proposed model of differential cell death in sexually dimorphic cells, with females experiencing more late-born cell death and males experiencing more early-born cell death. (**I**) Our web tool, available at flycns.com, provides an interactive interface for data exploration. Shown are examples of the interface “Expression” tab displaying comparative tSNEs illustrating the expression of multiple genes (*VAChT*, *VGlut*, *Gad1*, *Imp*, and *dati*) across *fru*+ neuronal cell types in the central brain.

To investigate this relationship across the entire central brain, we visualized sex differences in cell abundance, sexually dimorphic (*dsx*+ or *fru*+) versus monomorphic (*dsx*- and *fru*-) cell types, and *Imp* and *dati* expression (Figure 7D-F). Once again, we found a consistent correlation between sex-biased neuronal populations and birth-order (Figure 7G). Early-born neurons were overrepresented among female sexually dimorphic neurons compared to male dimorphic neurons (Figure 7G, upper left), while no sex difference in early vs. late markers was observed among monomorphic populations (Figure 7G, lower right). Comparing within-sex and across sexually dimorphic versus monomorphic neurons, revealed that female dimorphic neurons have an overrepresentation of early-born neurons relative to monomorphic female neurons (Figure 7G, upper right). Conversely, male dimorphic neurons have an underrepresentation of early-born neurons compared to male monomorphic neurons (Figure 7G, lower left).

These findings support a model where sexually dimorphic neurons arise through birth-order-dependent PCD within shared hemilineages (Figure 7H). In this framework, female- and male-specific neurons represent developmentally distinct subpopulations within a hemilineage, defined by sex-specific survival rather than by the generation of entirely distinct cell types. Thus, sexually dimorphic neurons – including the pC1 cluster – may be more accurately described as “paralogues” between the sexes rather than “orthologues”, arising from common progenitors but diverging via sex-biased elimination. Given that birth order is second only to hemilineage as a determinant of neuronal identity in the central brain, small sex-specific modifications in neuronal survival within a hemilineage lead to profound differences in circuit architecture and behaviour.

## DISCUSSION

It is widely assumed that the female and male nervous systems are composed of equivalent underlying cell types, with sex-specific features arising from differences in wiring, abundance, or physiology. Our findings challenge this view. By analysing sexed single-cell transcriptomes of the adult *Drosophila* central brain, we show that sexual dimorphism largely arises via the selective survival of distinct subpopulations of neurons within shared developmental lineages. By integrating transcriptional identity, sex, and markers of developmental histories, we provide a novel understanding of how behavioural diversity arises from a common developmental framework.

We took a data-driven approach to identify where sex influences neuronal identity across the adult central brain. Overall, sex-biased gene expression is limited, as only ∼1% of genes differ significantly between females and males, and most of these differences are modest in magnitude. However, when focusing on neuronal cell types showing the most pronounced differences in gene expression and cell abundance, we found that sexually dimorphic populations are consistently marked by expression of the sex determination genes *dsx* and *fru*. Gratifyingly, the convergence between our unbiased transcriptomic analysis and historical forward genetic screens for sex-specific behaviours, which identified both *dsx* and *fru*, validates both approaches and highlights the lasting insights offered by classical genetic approaches in behavioural neuroscience^68,69^.

Among the SDH genes, only *fru* and *dsx* show restricted, cell type-specific expression across the central brain, while upstream regulators like *Sxl* and *tra* are ubiquitously expressed. This finding suggests that the nervous system is transcriptionally poised for sexual differentiation but that only select subsets of neurons implement this program via sex-specific transcriptional networks. Intriguingly, *dsx* expression is always accompanied by *fru* expression within the same hemilineage, with subsets of cells co-expressing both. Prior work has shown that Dsx and Fru may co-regulate downstream targets and even bind regulatory elements of other SDH genes, suggesting a combinatorial code that stabilizes sexual identity^70,71^. This modular control, consistent with the theory of facilitated variation^72^, enables small regulatory shifts in expression timing or pattern to yield large-scale effects on behaviour.

Although the adult nervous system is generated through neurogenesis, it is sculpted by apoptosis, a mechanism used throughout the animal kingdom to generate alternative forms and functions^73^.

Neurons born at different temporal windows face different survival outcomes in both sexes. We observe a striking sex bias in neuronal survival based on birth order, with female-biased neurons enriched among early-born subtypes and male-biased neurons among late-born ones (Figure 7). Rather than both sexes retaining and modifying equivalent neurons, our data show that females and males retain different subsets of neurons from the same hemilineage, shaped by sex-specific patterns of Dsx- or Fru-dependent cell death (Figure 6). Fru is known to protect neurons from apoptosis in males^36,37^, while Dsx has been shown to both inhibit and promote apoptosis, depending on the cellular context^10,39,74–84^. A novel finding in our study is the extent of Fru-dependent cell death, resulting in early-born neurons being retained in females but eliminated in males (Figure 7). Although Fru in males has been shown to induce PCD^79^, this remains an underappreciated mode of creating dimorphism. Future studies following the development of this circuitry will shed further light on when and where these mechanisms are differentially used within each hemilineage.

Our findings support a model in which sex-specific neuronal diversity arises through selective modulation of shared developmental programs. The interplay between temporal identity and sex-specific factors shapes dimorphic outcomes, aligning with the principle that core developmental processes balance robustness and flexibility to allow neural circuits to adapt without compromising their fundamental structure^85,86^. In this framework, early-born, female-biased pathfinding neurons provide a stable scaffold, while late-born, male-biased neurons offer a flexible substrate for behavioural elaboration. This strategy of leveraging developmental pathways to selectively retain or prune temporal windows between the sexes may prove to be a model of exaptation^87^, whereby a trait adapted for one purpose, neurodevelopment, is co-opted for another, sexual differentiation, and may facilitate the evolution of dimorphic behaviours.

Our findings recast sexual differentiation in the nervous system as a developmentally embedded, resource-efficient process. Rather than constructing de novo sex-specific architectures, the brain leverages conserved spatial and temporal logic, tuning outcomes via selective modulation of cell survival. This principle may generalize across species. In vertebrates, although hormonal cues dominate models of sexual differentiation, increasing evidence points to conserved developmental mechanisms – e.g., differential neurogenesis, maturation, programmed cell death, and synaptogenesis – as key contributors to behavioural dimorphism^88,89^. As in *Drosophila*, these processes likely act on shared developmental templates, modified by sex at key regulatory nodes. Our study also acts as a powerful roadmap for future cross-species comparisons. We have shown lineage diversification without duplication, rather than evolving a new “female version” and “male version” of a circuit, a single hemilineage gives rise to both, with sex determining which neurons survive. This process is a resource-efficient evolutionary solution not limited to sexual differentiation and has also been noted in adaptive differences between species^90^. Implicit in this model is that neither the female nor the male brain represents the “default”, but rather both are sculpted from a shared “template”, resulting in two separate and distinct trajectories.

We have created a user-friendly web portal (https://www.flycns.com) to improve accessibility and usability for the wider community. This portal offers interactive web-based visualization of the atlases referenced in this study (Figure 7I). We have shown that our atlas achieves the resolution necessary to detect subtle transcriptional differences in rare cell types by dramatically expanding the number of cells sampled from the adult brain. Importantly, we show that many sexually dimorphic neurons arise from different developmental windows within the same lineage and are not one-to-one correlates of each other. This finding suggests that sex-specific transcriptional differences often reflect divergent developmental histories, underscoring the need for future single-cell analyses encompassing a broad developmental window.

---

**Figure S1.**
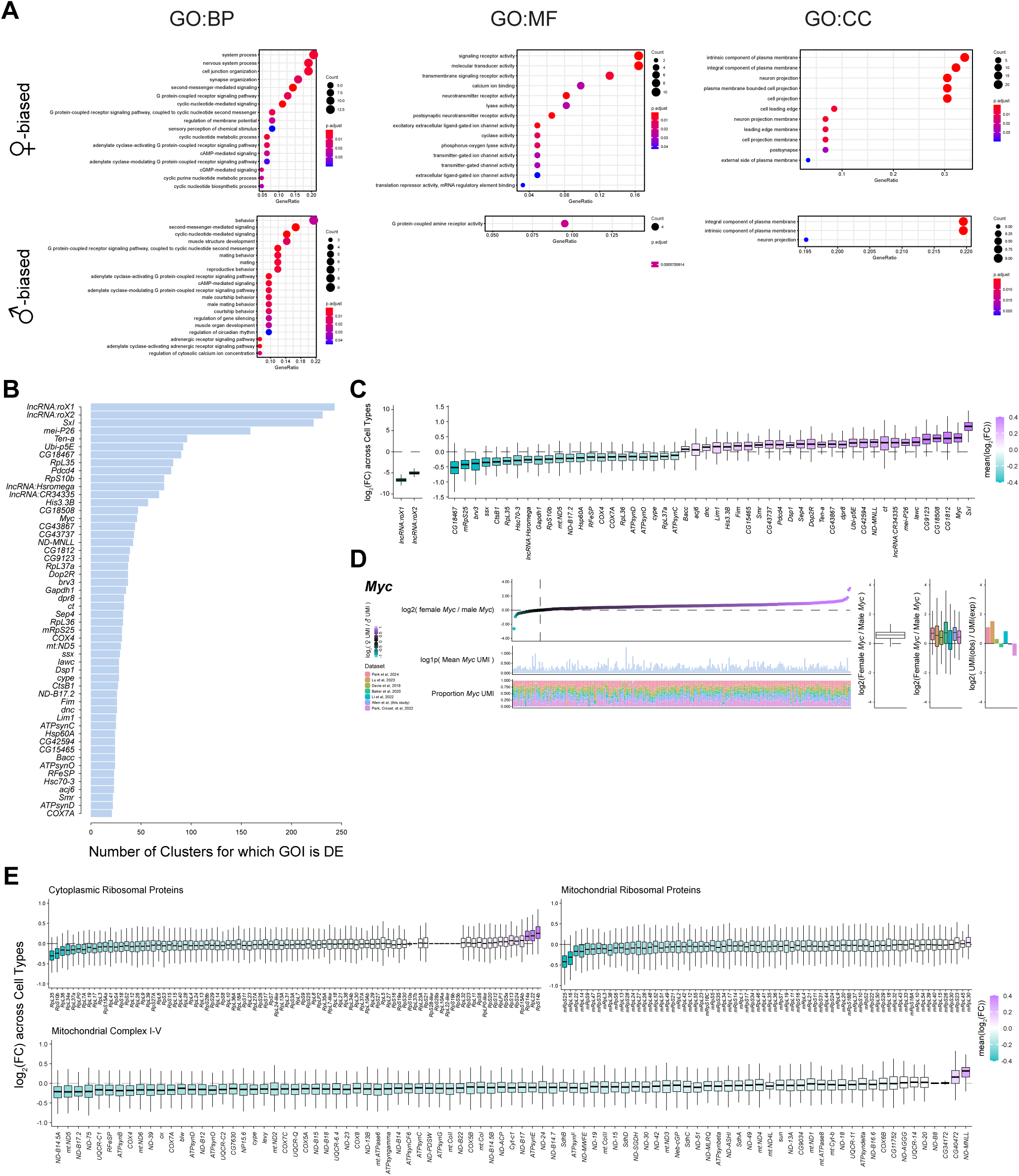
Differential expression across the brain. Related to Figure 1. (**A**) Gene ontology (GO) enrichment analysis of sex-biased genes in female (top) and male (bottom) central brain neurons. GO enrichment for biological processes (left), molecular function (middle) and cellular compartments (right) shown. (**B**) Bar graph displaying the number of neuronal cell types (clusters) with differential expression (DE) of gene of interest (GOI) with p_adj_-value < 0.01 and any log_2_ fold change value. (**C**) Boxplots showing log_2_ fold-change of sex-biased gene expression between females and males across all cell types in the central brain. (**D**) *Myc* expression is ubiquitously female-biased. Left: *Myc* expression values (female/male) (top), expression levels (mean UMI) and proportion detected in each dataset (proportion UMI) across all neuronal types in the central brain. Right: box plots of *Myc* expression across all datasets (left), individual datasets (middle), and bar graphs showing deviations from expected expression levels (observed/expected UMI). (**E**) Boxplots showing log_2_ fold-change in expression between the sexes of cytoplasmic (left) and mitochondrial (right) ribosomal protein genes (top), and mitochondrial complex I-V genes (bottom) across all neuronal types in the central brain.

**Figure S2.**
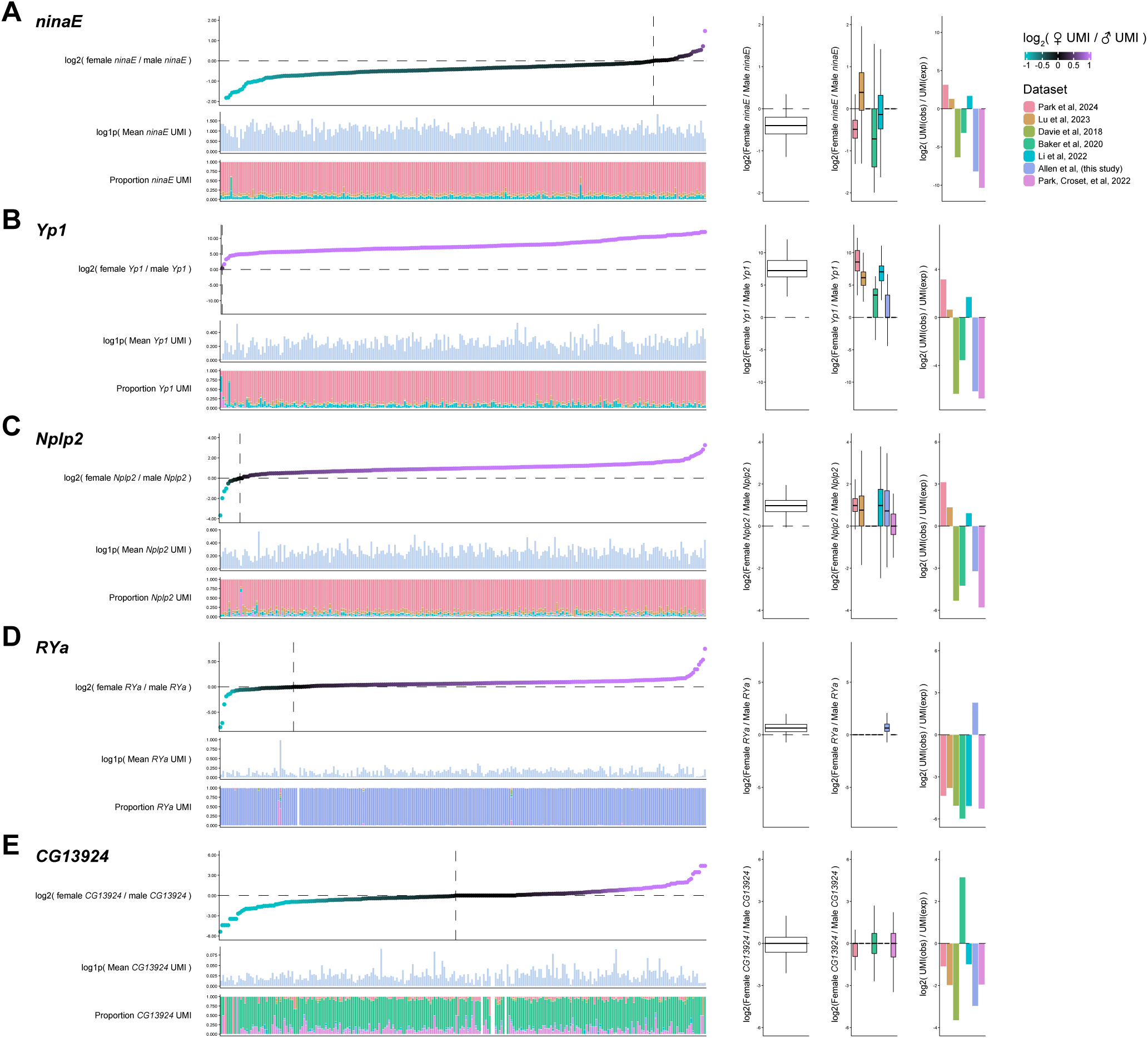
Identification of dataset-specific artefactually sex-biased genes. Related to Figure 1. (**A**-**E**) Left: GOI expression values (female/male) (top), expression levels (mean UMI) and proportion detected in each dataset (proportion UMI) across all neuronal types in the central brain. Right: box plots of GOI expression across all datasets (left), individual datasets (middle) and bar graphs showing deviations from expected expression levels (observed/expected UMI). Genes: *neither inactivation nor afterpotential E* (*ninaE*, **A**), *Yolk protein 1*(*Yp1*, **B**), *Neuropeptide-like precursor 2* (*Nplp2*, **C**), *RYamide* (*RYa*, **D**), *CG13924* (**E**).

**Figure S3.**
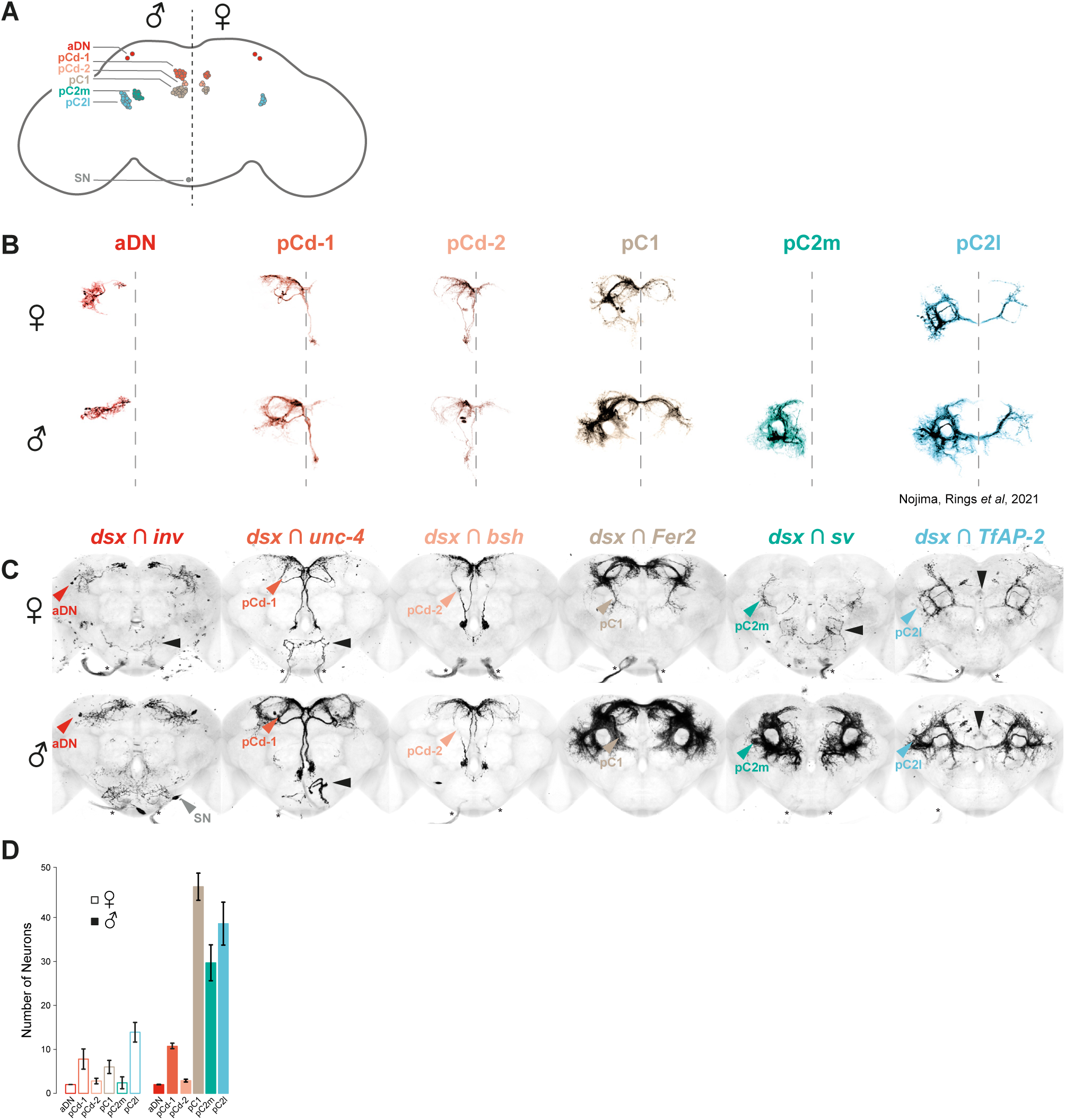
*dsx+* neuroanatomy. Related to Figure 2. (**A**) A schematic drawing of *dsx*+ neuronal subtypes in the male (left) and female (right) brain. (**B**) Previously published single clone MARCM results of *dsx*+ neuronal populations (Nojima et al., 2021^32^). (**C**) Whole-mount immunofluorescence of central brains with *dsx^DBD^* and *TF^AD^*hemi-driver combinations uniquely labels predicted *dsx*+ neuronal populations. Images registered onto a template brain; arrowheads indicate peripheral sensory neuron innervations, and asterisks indicate ectopic expression of the reporter. (**D**). Cell counts of genetic intersections recover similar numbers to previous MARCM results (as shown in B).

**Figure S4.**
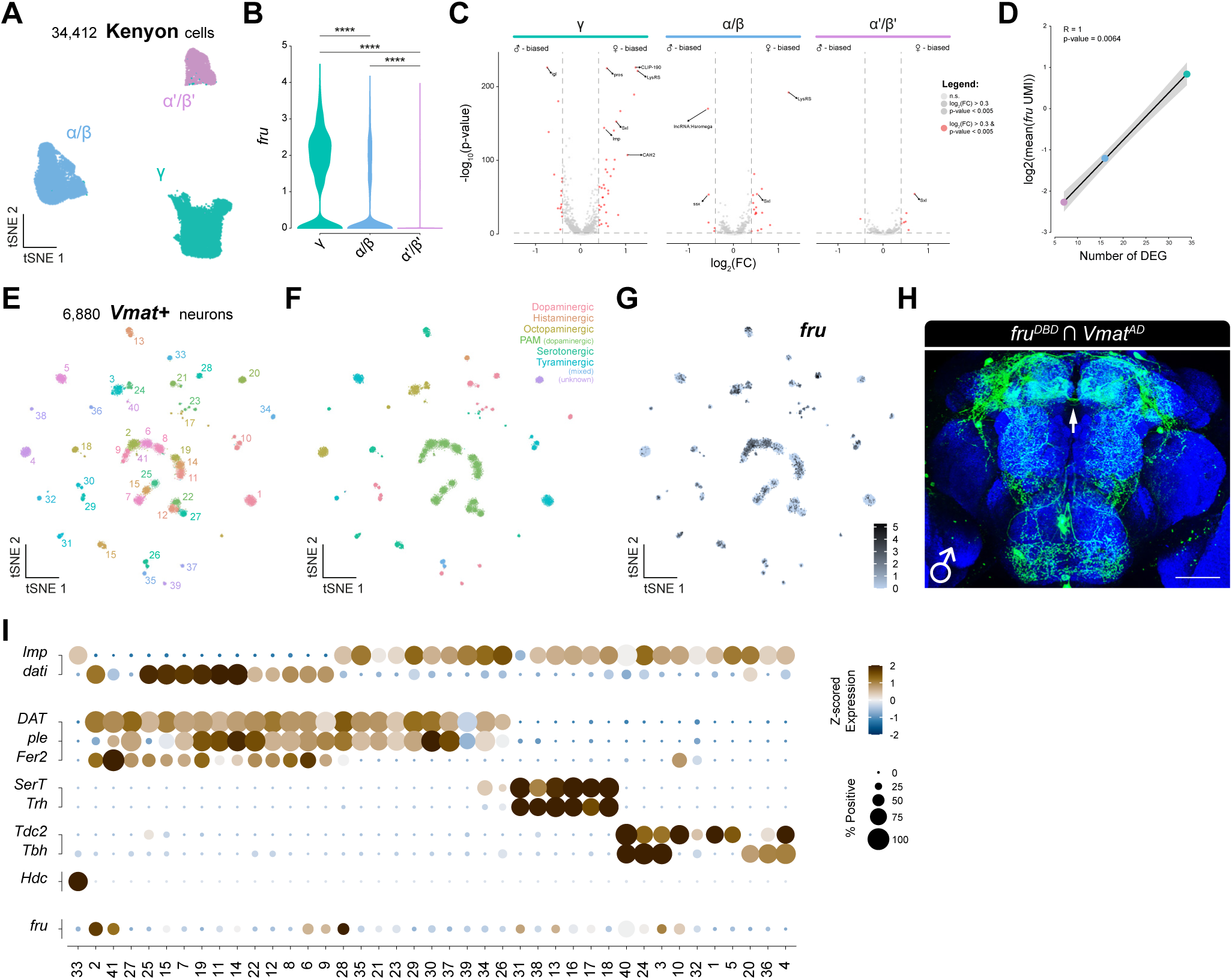
*fruitless* expression in Kenyon cells and monoaminergic neurons. Related to Figure 3. (**A**) tSNE of subclustered Kenyon cells. (**B**) Violin plot of *fru* expression in the 3 Kenyon cell subtypes. ****, p_adj_-value < 0.0001 (Wilcoxon Signed-Rank Test, Bonferroni corrected). (**C**) Volcano plot of differentially expressed genes across the 3 Kenyon cell subtypes. (**D**) Correlation of number of differentially expressed genes (abs(log_2_(FC)) > 0.4, p_adj_-value < 0.05) with *fru* expression across the 3 Kenyon cell subtypes. (**E**) tSNE of sub-clustered monoaminergic neurons (see^19^) coloured by unsupervised Leiden clustering. (**F**) tSNE of sub-clustered monoaminergic neurons (see^19^) coloured by broad monoamine usage. (**G**) tSNE of *fru* expression in sub-clustered monoaminergic neurons. (**H**) Registered whole-mount immunofluorescence of male central brain with *fru^DBD^* and *Vmat^AD^* intersection labelling *fru*-expressing monoaminergic neurons. Arrow indicating PAM-associated midline crossing. (**I**) Dot plot of birth order genes (*Imp, dati*), monoaminergic subtype markers (*DAT, ple, Fer2, SerT, Trh, Tdc2, Tbh,* and *Hdc*), and *fru* across all monoaminergic cell types.

**Figure S5.**
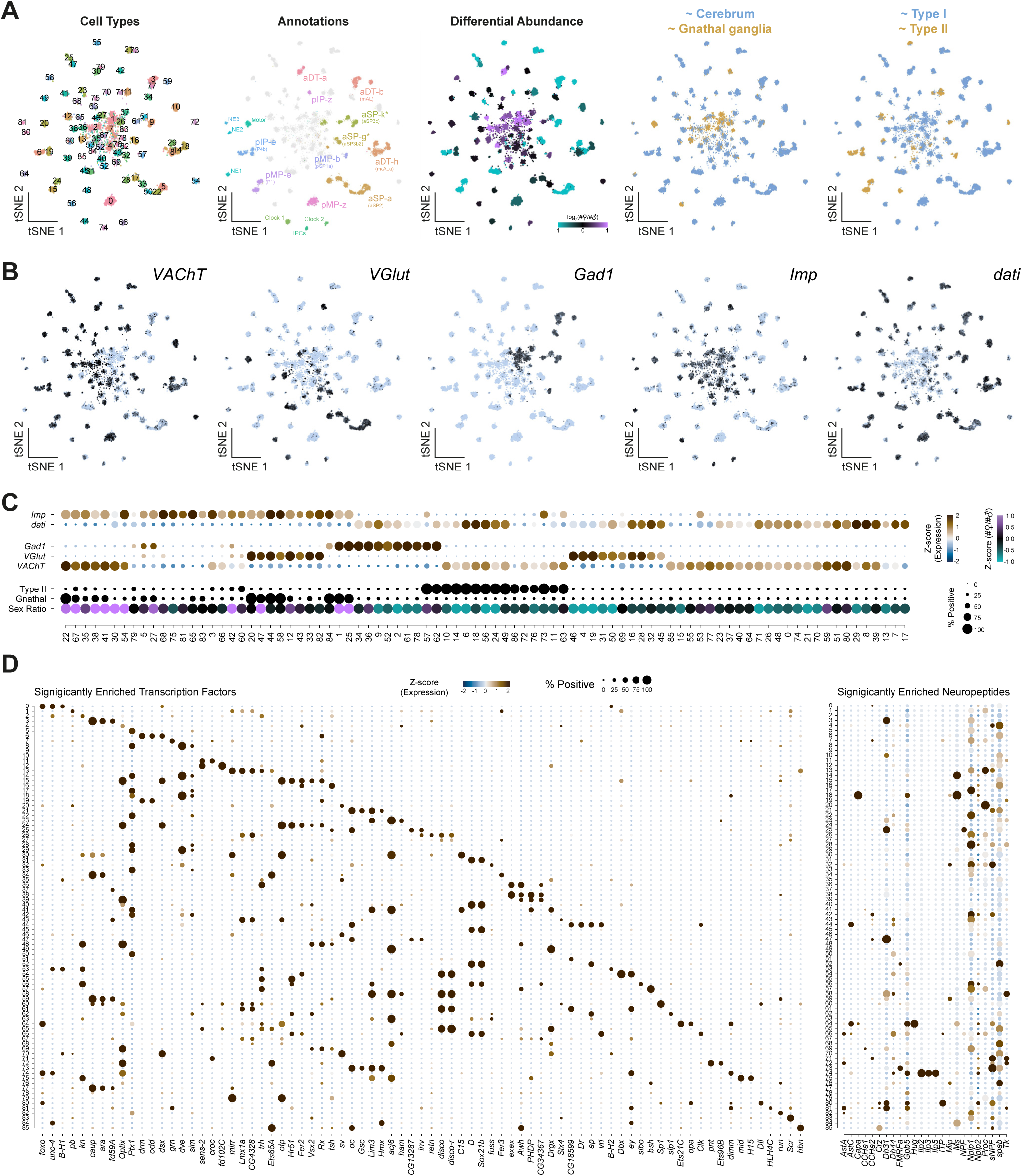
Organisational principles of the *fru*+ central brain neuron atlas. Related to Figure 3. (**A**) tSNEs of *fru*+ atlas showing transcriptionally distinct cell types (by colour), annotated cell types, differential abundance (female/male), neuromere (cerebrum vs. gnathal ganglia) and neuroblast lineage type (type I vs. type II). (**B**) tSNEs of *fru*+ atlas showing neurotransmitter usage (*VAChT*, *VGlut*, *Gad1*), and early (*Imp*) vs. late (*dati*) birth order. (**C**) Dot plot showing the expression levels of key biomarkers for birth order (*Imp* and *dati*), neurotransmitter usage (*VAChT*, *VGlut*, *Gad1*), type II derived neuroblast lineage, gnathal ganglia derived and sex-biased cell abundance across *fru+* neuronal subtypes. (**D**) Dot plot of the expression of significantly enriched transcription factors (left) and neuropeptides (right) across *fru+* neuronal subtypes.

**Figure S6.**
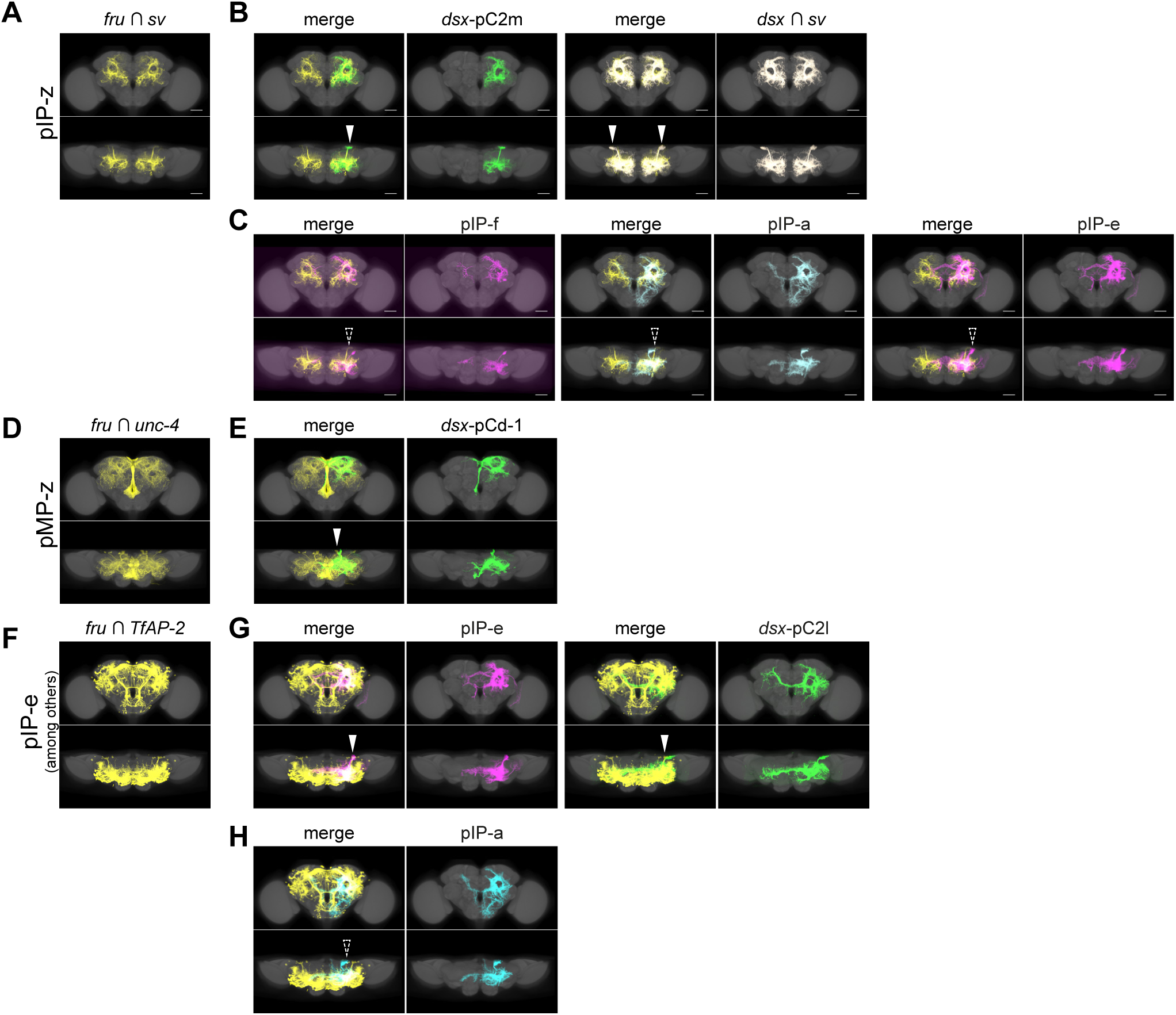
Neuroanatomical validation of *fru+* annotations. Related to Figure 3. (**A**) Registered whole-mount immunofluorescence of male central brains with *fru^DBD^* and *sv^AD^* intersection labelling what we are calling pIP-Z. (**B**) Registered whole-mount immunofluorescence of male central brains with *fru^DBD^*/*sv^AD^* intersection, *dsx+* pC2m MARCM clone (originally from^32^), and with *dsx^DBD^*/*sv^AD^* intersection showing overlapping expression in hemilineage-associated tracks (solid arrowheads). (**C**) Registered whole-mount immunofluorescence of male central brains with *fru^DBD^*/*sv^AD^* intersection and *fru+* MARCM clones pIP-f, pIP-a, and pIP-e (originally from^34^) showing lack of overlapping expression in hemilineage-associated tracks (dashed arrowheads). (**D**) Registered whole-mount immunofluorescence of central brains with *fru^DBD^* and unc-4*^AD^* intersection labelling what we are calling pMP-Z. (**E**) Registered whole-mount immunofluorescence of central brains with *fru^DBD^*/*unc-4^AD^* intersection and *dsx+* pCd-1 MARCM clone showing overlapping expression in hemilineage-associated tracks (solid arrowhead). (**F**) Registered whole-mount immunofluorescence of central brains with *fru^DBD^* and *TfAP-2^AD^* intersection labelling pIP-e, as well as multiple other cell types. (**G**) Registered whole-mount immunofluorescence of central brains with *fru^DBD^*/*TfAP-2^AD^* intersection, *fru+* pIP-e MARCM clone (originally from^34^), *dsx+* pC2l MARCM clone (originally from^32^), showing overlapping expression in hemilineage-associated tracks (solid arrowheads). (**H**) Registered whole-mount immunofluorescence of central brains with *fru^DBD^*/*TfAP-2^AD^* intersection and *fru+* pIP-a MARCM clone (originally from^34^) showing lack of overlapping expression in hemilineage-associated tracks (dashed arrowhead). Previous data suggested that *fru+* pIP-a is derived from the DL2 lineage along with *dsx+* pC2l^37^. These data suggest that, in fact, it is *fru+* pIP-e that is associated with *dsx+* pC2l and the DL2 lineage and not pIP-a. For all panels, frontal view (top) and horizontal view (bottom) are shown.

**Figure S7.**
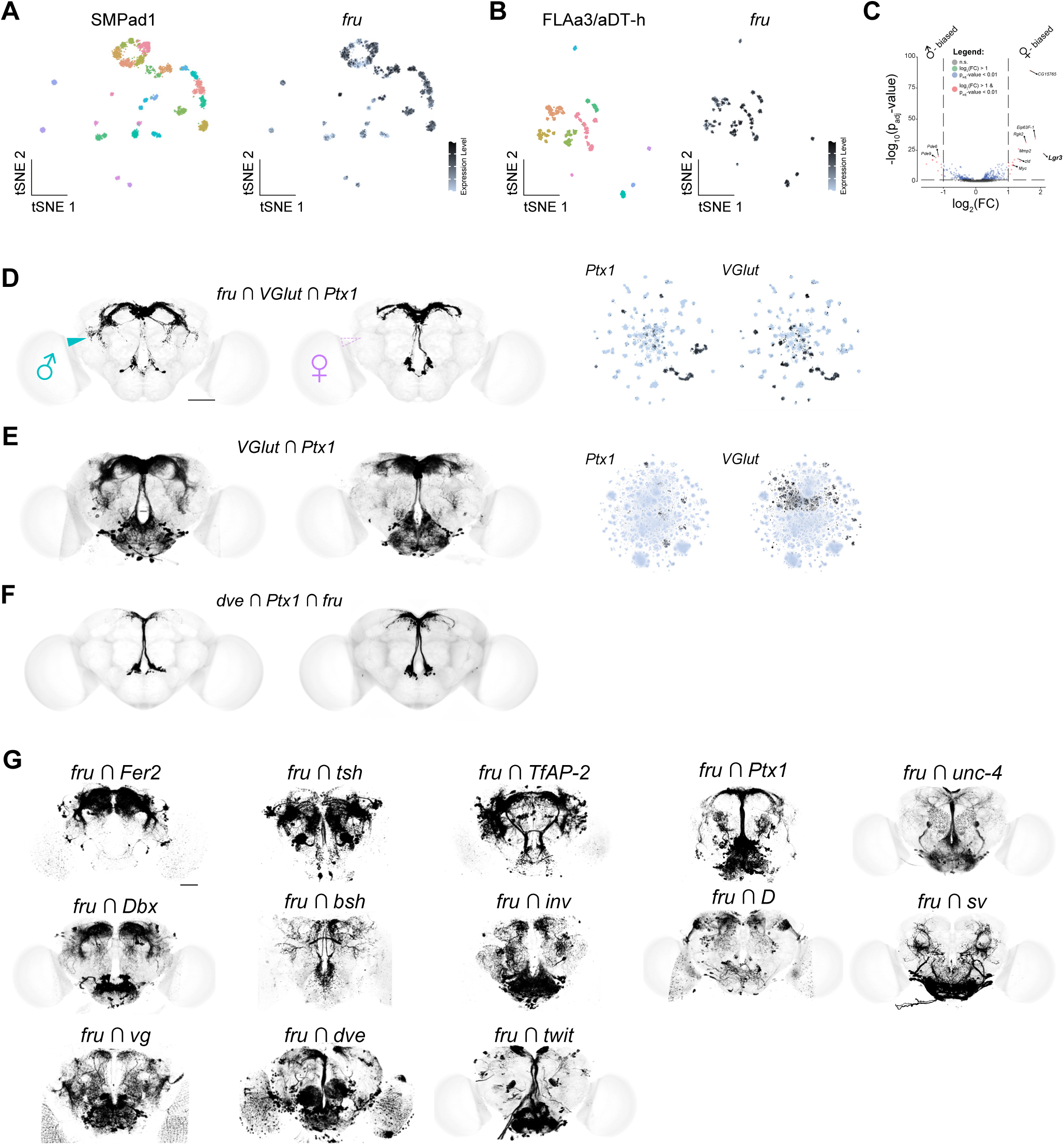
*fru* intersections. Related to Figure 3. (**A**) t-SNE of subclustered SMPad1 (left) and *fru* expression in SMPad1 (right). Most SMPad1 subtypes express *fru*. (**B**) t-SNE of subclustered FLAa3 (left) and *fru* expression in FLAa3 (right). All FLAa3 subtypes express *fru* and so FLAa3 and *fru+* aDT-h are synonymous. (**C**) Volcano plot of differentially expressed genes in FLAa3/aDT-h. (**D**) Registered, segmented, whole-mount immunofluorescence triple genetic intersection of *fru*, *Ptx1*, and *VGlut*, labelling SMPad1 hemilineage with arrowheads indicating male-specific neurites (left). tSNEs of *Ptx1* and *VGlut* in the *fru+* atlas (right). (**E**) Registered, unsegmented, whole-mount immunofluorescence of central brains with *Ptx1* and *VGlut* intersection labelling SMPad1 hemilineage, as well as multiple other cell types (left). tSNEs of *Ptx1* and *VGlut* in the full central brain neuronal atlas (right). (**F**) Registered, unsegmented, whole-mount immunofluorescence of central brains with *dve*, *Ptx1*, and *fru*. (**G**) Multiple attempts at genetic intersection to annotate *fru+* transcriptional clusters consistently labelled many cell types often obscuring identification. Registered whole-mount immunofluorescence of male central brains with *fru* intersected with: *Fer2*, *tsh*, *TfAP-2*, *Ptx1*, *unc-4*, *Dbx*, *bsh, inv*, *D*, *sv*, *vg*, *dve*, and *twit*.

**Figure S8.**
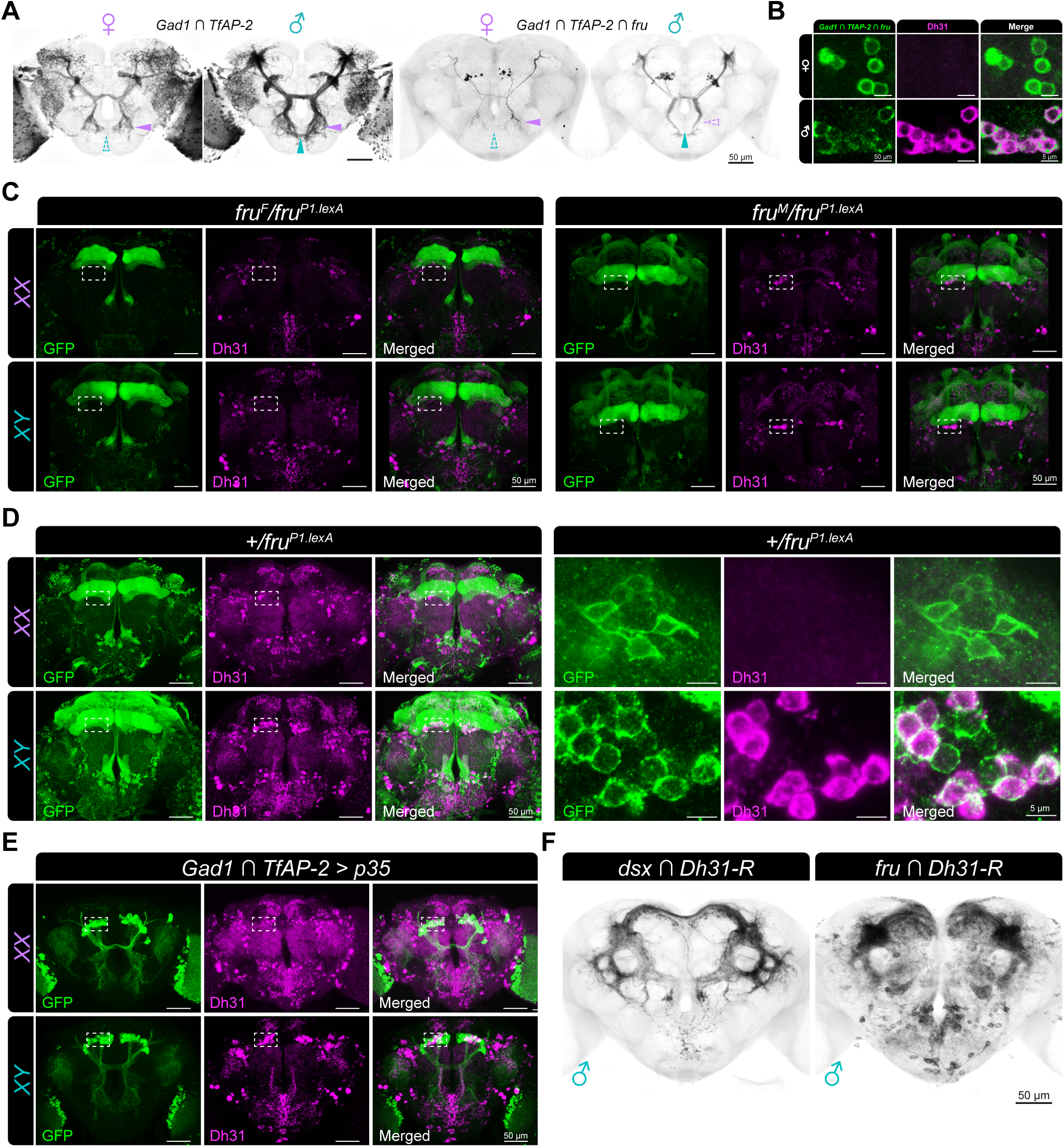
Male-specific expression of the Dh31 in mAL neurons. Related to Figure 4. (**A**) Genetic intersection of CREa1 ventral hemilineage and mAL neurons. Unsegmented immunofluorescence images of *Gad1 ∩ TfAP-2* (left) and *Gad1 ∩ TfAP-2 ∩ fru* (right) central brains from Figure 4E and 4F, respectively. Arrowheads highlight the anatomical presence (solid) or absence (dashed) of neurite projections between the sexes. (**B**) Whole mount immunofluorescence closeup images of individual section of soma in *Gad1* ∩ *TfAP-2 ∩ fru* intersection expressing mGFP (green) and counterstained with anti-Dh31 (magenta) and merged (white). Female (top) and male (bottom). (**C**) Whole brain immunofluorescence images of Dh31 expression in *fru*+ neurons in hemizygous *fru^F^* females (*fru^F^*/*fru^P^*^1^*^.LexA^; left)* and males (*fru^M^*/*fru^P^*^1^*^.LexA^; right*). *fru^P^*^1^*^.LexA^*driven mGFP expression (green) shown, counterstained with anti-Dh31 (magenta) and merged (white). Dashed white box encompasses region highlighted in closeup individual sections of soma shown on in Figure 4I. (**D**) Whole brain immunofluorescence images of Dh31 expression in the control hemizygous *fru*-null (*+*/*fru^P^*^1^*^.LexA^*) females (XX, top) and males (XY, bottom). Whole brains shown on left, with dashed white box encompasses region highlighted in closeup individual sections of soma shown on the right. (**E**) Whole brain immunofluorescence images of Dh31 expression in p35 expressing *Gad1* ∩ *TfAP-2* intersections expressing mGFP (green) and counterstained with anti-Dh31 (magenta) and merged (white). Female (top) and male (bottom) whole brains shown on left, with dashed white box encompasses region highlighted in closeup individual sections of soma shown on in Figure 4J. (**F**) *Dh31-R* expression in sexually dimorphic central brain neurons. Immunofluorescence images of *dsx ∩ Dh31-R* (left) and fru *∩ Dh31-R* (right) in male central brains.

**Figure S9.**
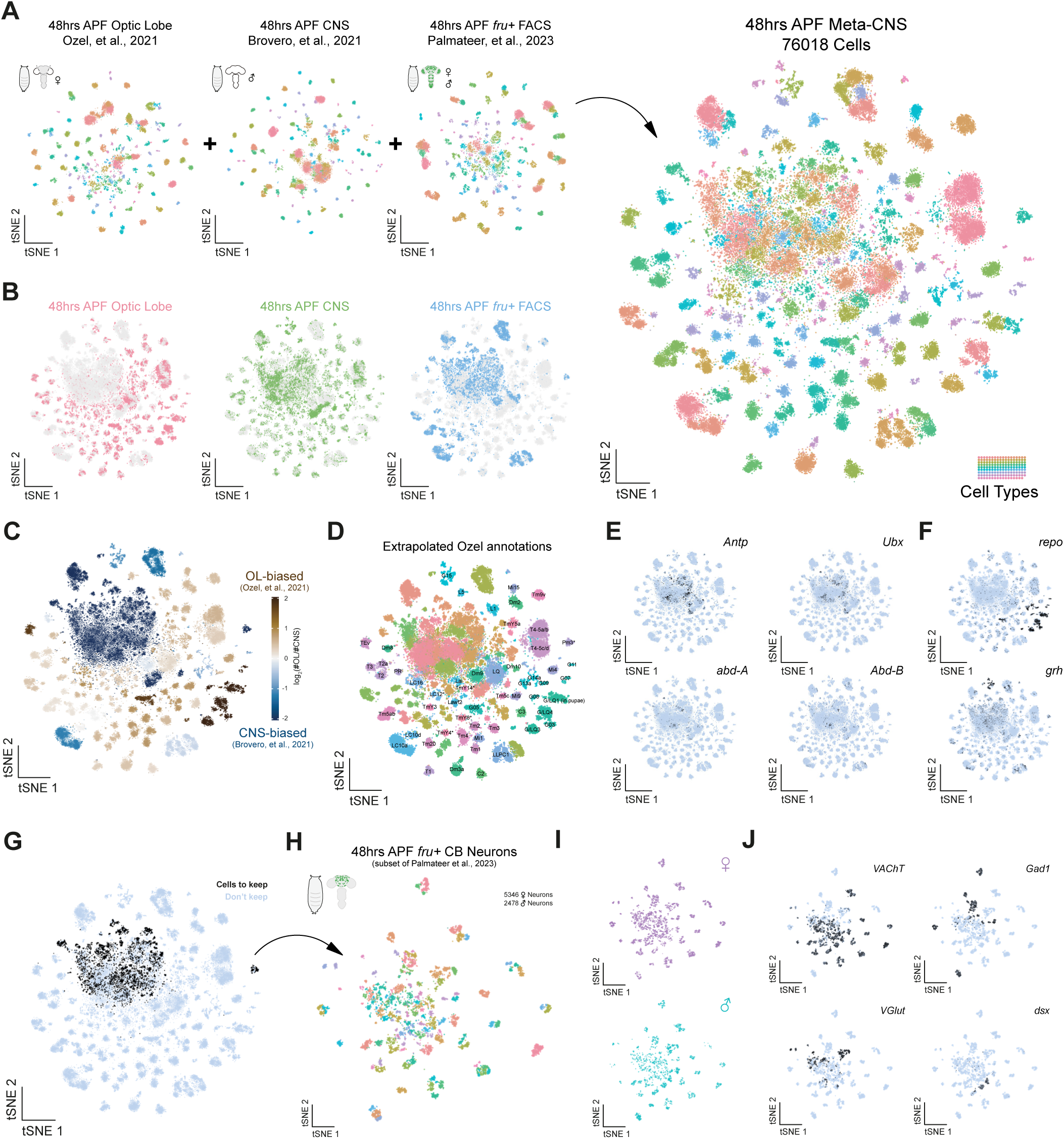
Re-analysis and annotation of *fru*+ 48hr APF meta-CNS atlas. Related to Figure 4. (**A**) Integration of three re-analysed 48hr after puparium formation (APF) scRNA-seq datasets (left): optic lobe^91^, CNS^92^, and *fru+* FAC-sorted neurons^52^, resulting in a 76,018-cell Meta-CNS atlas (right), coloured by transcriptionally defined cell types. (**B**) tSNEs showing dataset-specific contribution to the Meta-CNS atlas. (**C**) tSNE showing relative representation of the optic lobe (OL) and CNS datasets in the Meta-CNS atlas. (**D**) Cell type annotations from Ozel *et al.,* 2021^91^, extrapolated across the integrated atlas to infer transcriptionally distinct cell type identities. (**E**) tSNEs showing expression of homeotic genes (*Antp*, *Ubx*, *abd-A*, *Abd-B*) used to define VNC derived cells. (**F**) tSNEs showing expression of glial (*repo*) and epithelial (*grh*) marker genes used to define non-neuronal identities. (**G**) tSNE highlighting cells retained for *fru*+ central brain atlas based on quality control and annotation confidence. (**H**) 48hrs APF *fru*+ central brain neuron atlas coloured by transcriptionally defined cell types. (**I**) tSNEs showing female (top) and male (bottom) cells in 48hrs APF *fru*+ central brain neuron atlas. (**J**) tSNEs showing fast-acting neurotransmitter-defining gene expression (*VAChT*, *VGlut*, *Gad1*) in 48hrs APF *fru*+ central brain neuron atlas.

**Figure S10.**
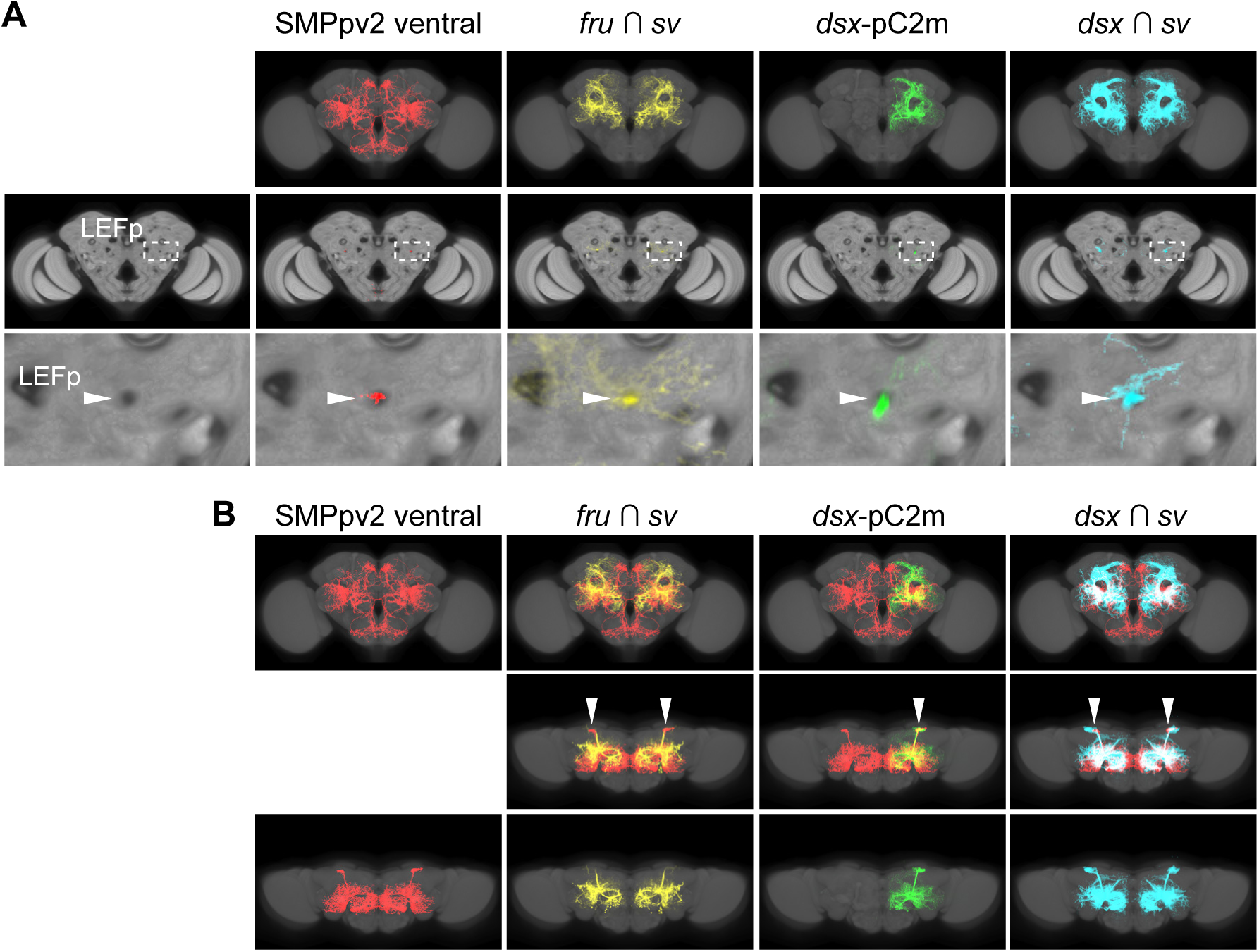
Identifying *dsx+* pC2m and *fru+* pIP-z as members of the hemilineage SMPpv2 ventral. Related to Figure 5. (**A**) Full Z-projection (top), single posterior slice (middle), and zoom-in (bottom) of (from left to right) SMPpv2 ventral skeletons (Flywire), pIP-z (*fru ∩ sv*), pC2m (*dsx^Gal^*^4^ MARCM clone), and pC2m (*dsx* ∩ *sv*). Dashed boxes and solid arrowheads highlight the hole in the anti-Brp signal corresponding to the BP-104 positive LEFp bundle. SMPpv2 ventral is the only hemilineage to contribute to this bundle^54,55^, thus *fru+* pIP-z and *dsx+* pC2m are members of the SMPpv2 ventral hemilineage. (**B**) Registered SMPpv2 ventral skeletons (Flywire) and whole-mount immunofluorescence of central brains with *fru^DBD^*/*sv^AD^*intersection, *dsx+* pC2m MARCM clone (originally from^32^), and with *dsx^DBD^*/*sv^AD^* intersection showing overlapping expression in SMPpv2 ventral hemilineage-associated tracks (solid arrowheads). Frontal view (top) and horizontal views (middle and bottom) are shown. Note, the Flywire SMPpv2 ventral skeleton is derived from a female brain, while all whole-mount immunofluorescence images are derived from male brains.

**Figure S11.**
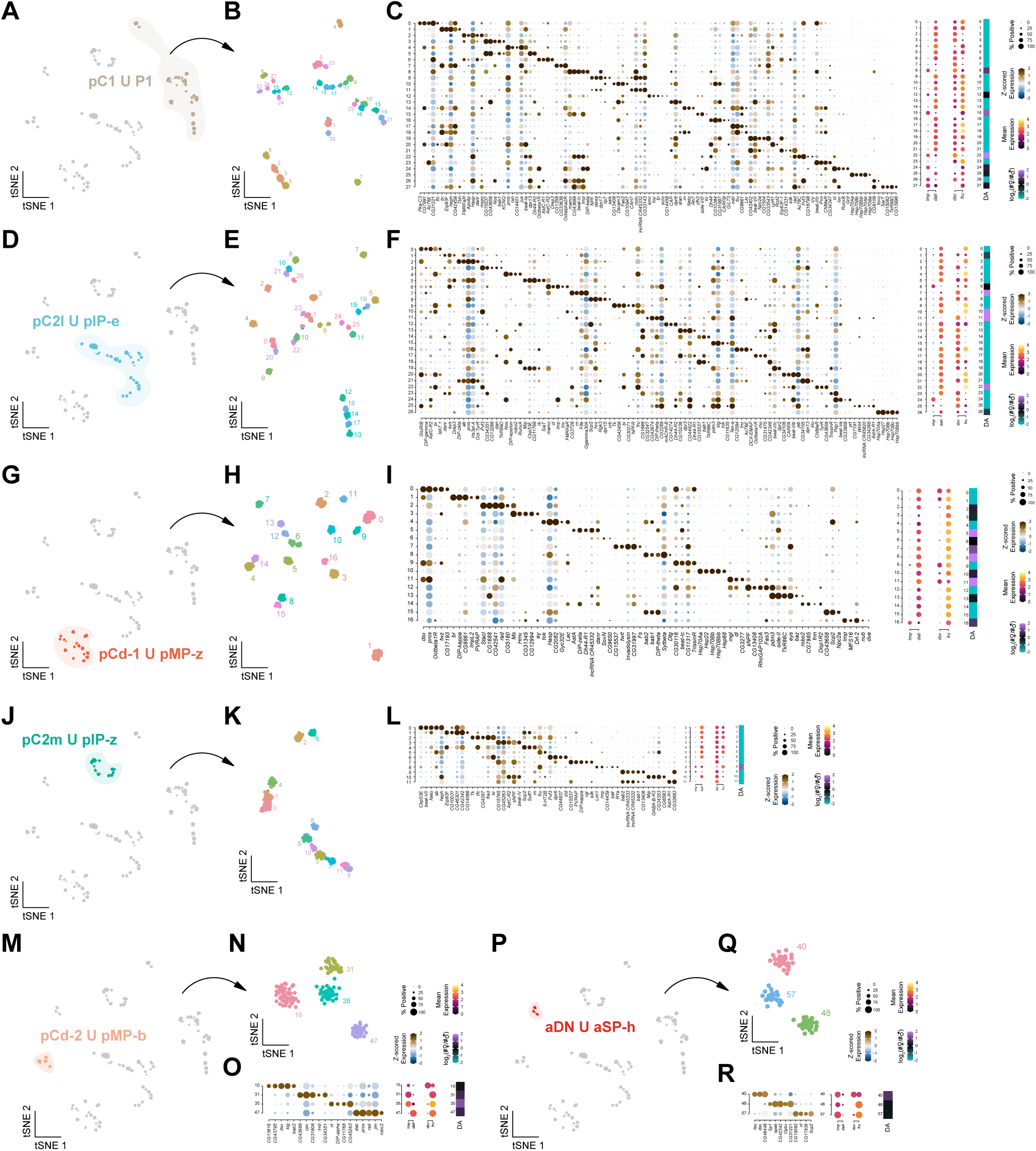
Identification of vast transcriptional heterogeneity amongst *dsx* and *fru* co-expressing neuronal populations. Related to Figure 5. (**A**) tSNE of subclustered *dsx*+ U *fru*+ neuronal cells with shared hemilineage identities highlighting the *dsx+* pC1 and *fru+* P1 (pMP-e) cells of DM4 dorsal. (**B**) tSNE of further subclustered pC1/P1 neurons, coloured by the 28 distinct cell subtypes. (**C**) Dot plots and heatmaps showing the significantly enriched genes (left), birth order genes (*Imp*, *dati*) and sex determination genes (*dsx*, *fru*) (middle), and differential abundance of cell numbers between the sexes (DA, right) across the 28 distinct cell subtypes of pC1/P1. (**D** tSNE of subclustered *dsx*+ U *fru*+ neuronal cells with shared hemilineage identities highlighting the *dsx+* pC2l and *fru+* pIP-e cells of DL2 ventral. (**E**) tSNE of further subclustered pC2l/pIP-e neurons, coloured by the 27 distinct cell subtypes. (**F**) Dot plots and heatmaps showing the significantly enriched genes (left), birth order genes (*Imp*, *dati*) and sex determination genes (*dsx*, *fru*) (middle), and differential abundance of cell numbers between the sexes (DA, right) across the 27 distinct cell subtypes of pC2l/pIP-e. (**G**) tSNE of subclustered *dsx*+ U *fru*+ neuronal cells with shared hemilineage identities highlighting the *dsx+* pCd-1 and *fru+* pMP-z cells of SMPpv1. (**H**) tSNE of further subclustered pCd-1/pMP-z neurons, coloured by the 17 distinct cell subtypes. (**I**) Dot plots and heatmaps showing the significantly enriched genes (left), birth order genes (*Imp*, *dati*) and sex determination genes (*dsx*, *fru*) (middle), and differential abundance of cell numbers between the sexes (DA, right) across the 17 distinct cell subtypes of pCd-1/pMP-z. (**J**) tSNE of subclustered *dsx*+ U *fru*+ neuronal cells with shared hemilineage identities highlighting the *dsx+* pC2m and *fru+* pIP-z cells of SMPpv2 ventral. (**K**) tSNE of further subclustered pC2m/pIP-z neurons, coloured by the 12 distinct cell subtypes. (**L**) Dot plots and heatmaps showing the significantly enriched genes (left), birth order genes (*Imp*, *dati*) and sex determination genes (*dsx*, *fru*) (middle), and differential abundance of cell numbers between the sexes (DA, right) across the 12 distinct cell subtypes of pC2m/pIP-z. (**M**) tSNE of subclustered *dsx*+ U *fru*+ neuronal cells with shared hemilineage identities highlighting the *dsx+* pCd-2 and *fru+* pMP-b cells of DM2 central. (**N**) tSNE of further subclustered pCd-2/pMP-b neurons, coloured by the 4 distinct cell subtypes. (**O**) Dot plots and heatmaps showing the significantly enriched genes (left), birth order genes (*Imp*, *dati*) and sex determination genes (*dsx*, *fru*) (middle), and differential abundance of cell numbers between the sexes (DA, right) across the 4 distinct cell subtypes of pCd-2/pMP-b. (**P**) tSNE of subclustered *dsx*+ U *fru*+ neuronal cells with shared hemilineage identities highlighting the *dsx+* aDN and *fru+* aSP-h cells of LHl2 dorsal. (**Q**) tSNE of further subclustered aDN/aSP-h neurons, coloured by the 3 distinct cell subtypes. (**R**) Dot plots and heatmaps showing the significantly enriched genes (left), birth order genes (*Imp*, *dati*) and sex determination genes (*dsx*, *fru*) (middle), and differential abundance of cell numbers between the sexes (DA, right) across the 3 distinct cell subtypes of aDN/aSP-h.

**Figure S12.**
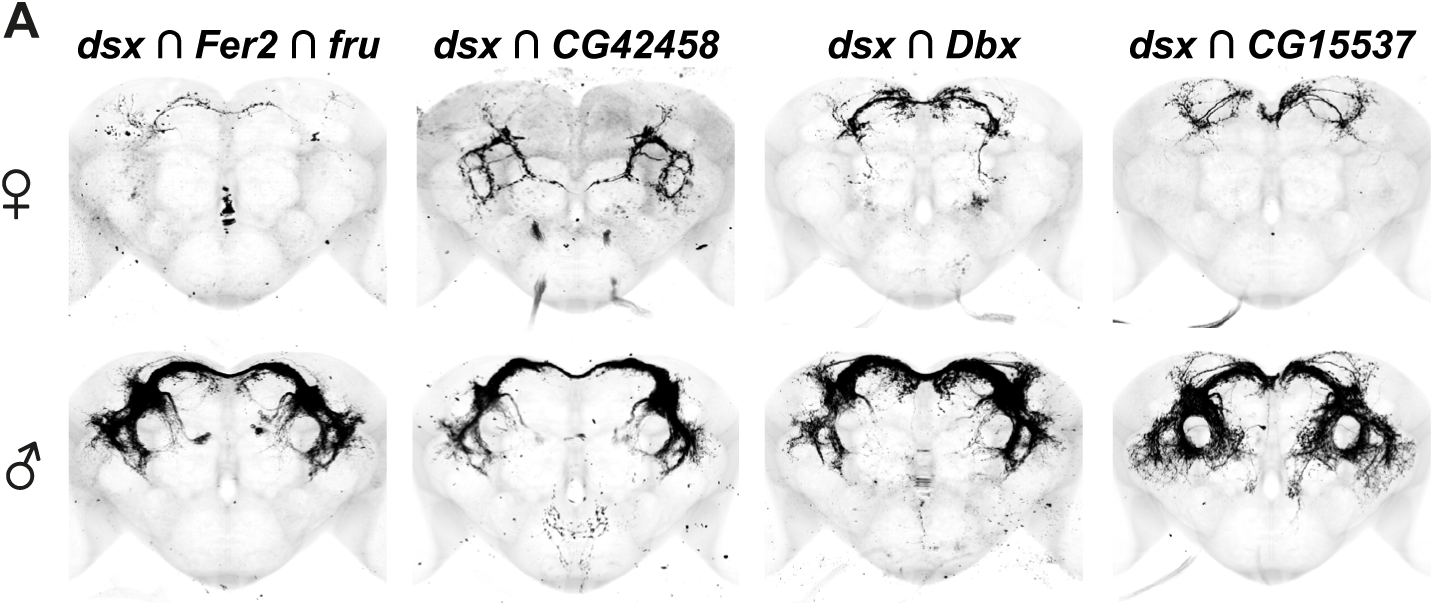
Genetic intersections of transcriptionally distinct pC1 subtypes. Related to Figure 6. (**A**). Unsegmented whole brain immunofluorescence light microscopy images in females (top) and males (bottom) from Figure 6D. Split-Gal4 driven mGFP expression shown in black, images co-registered onto a template brain.

## STAR METHODS

### KEY RESOURCES TABLE

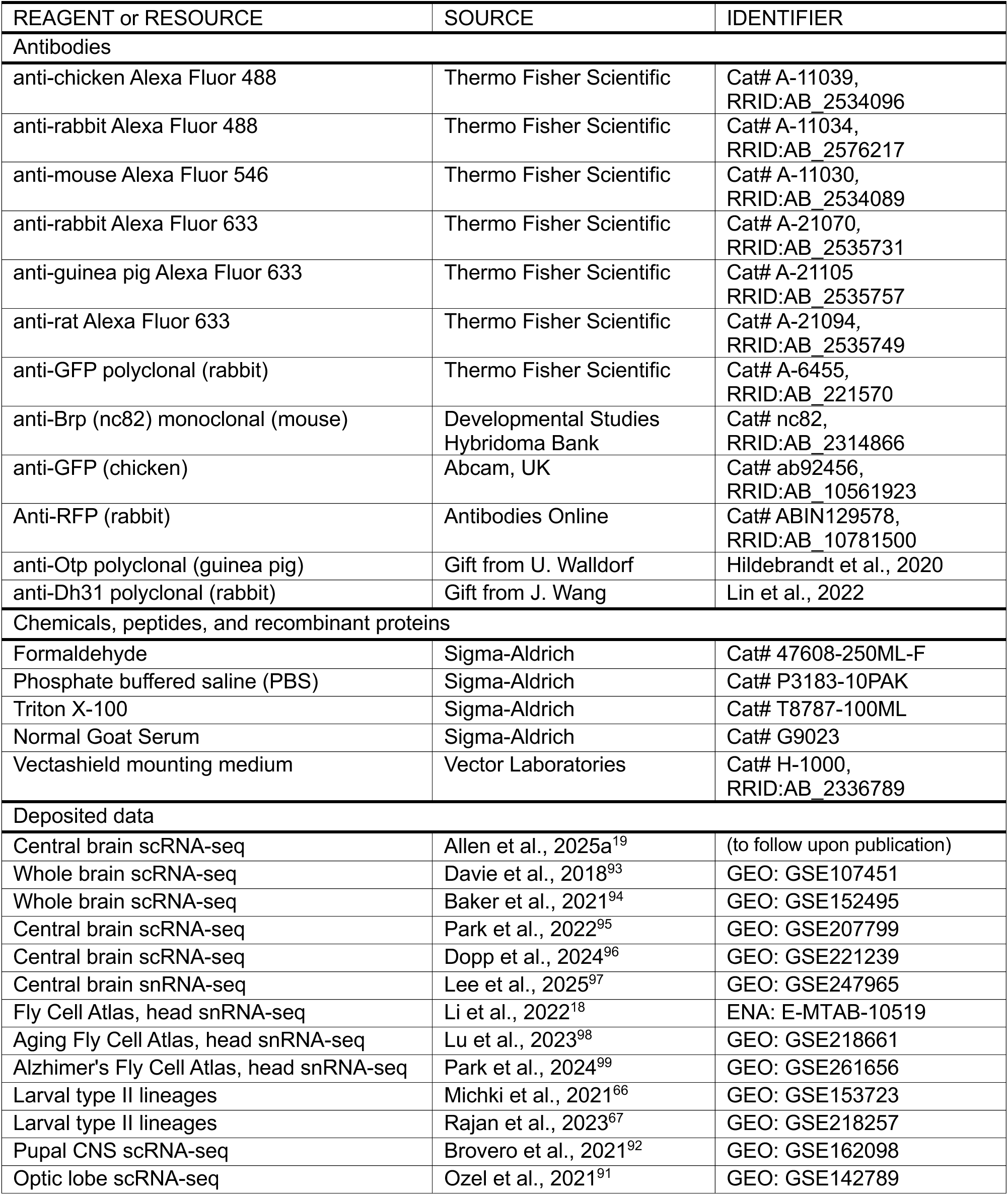
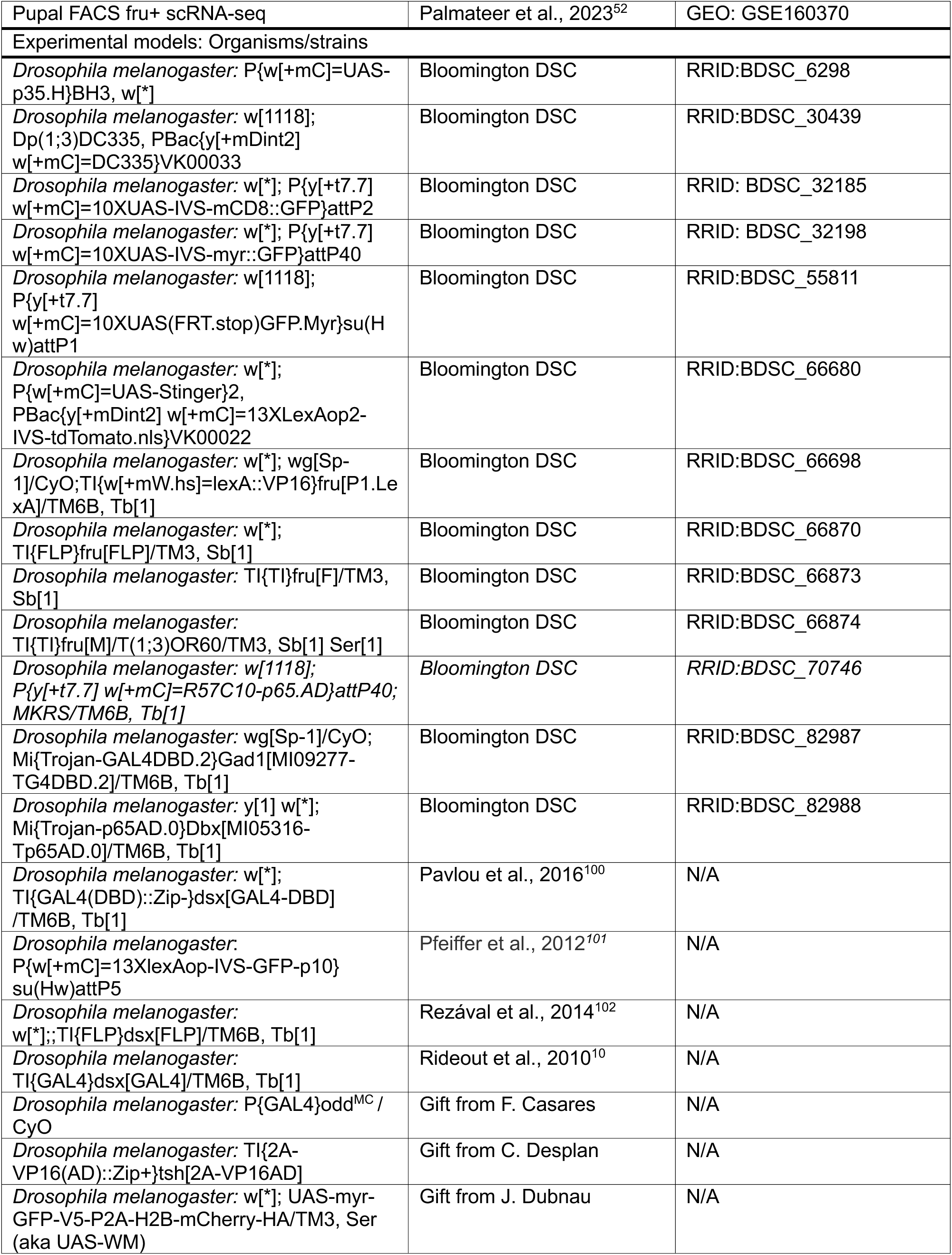
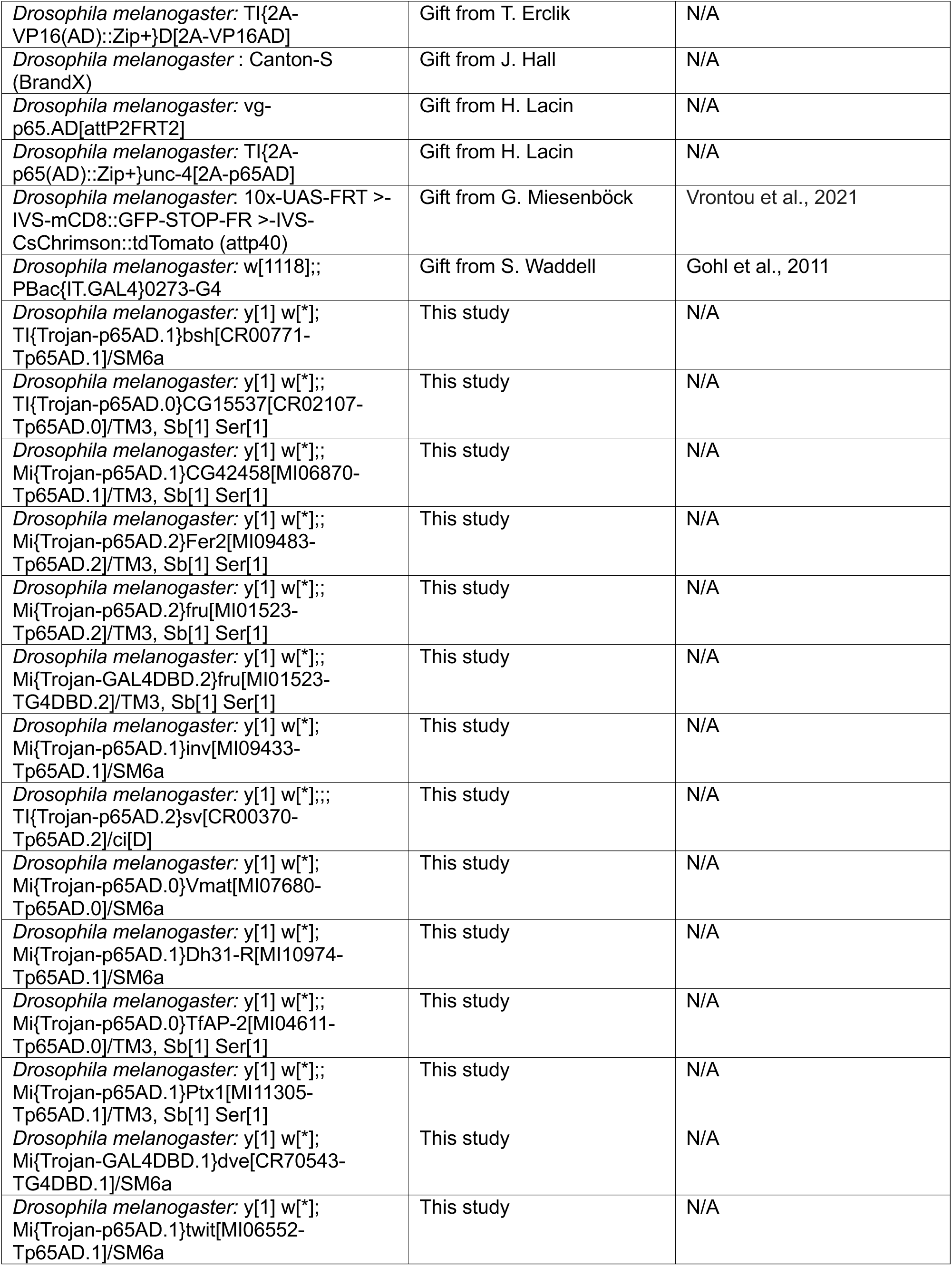
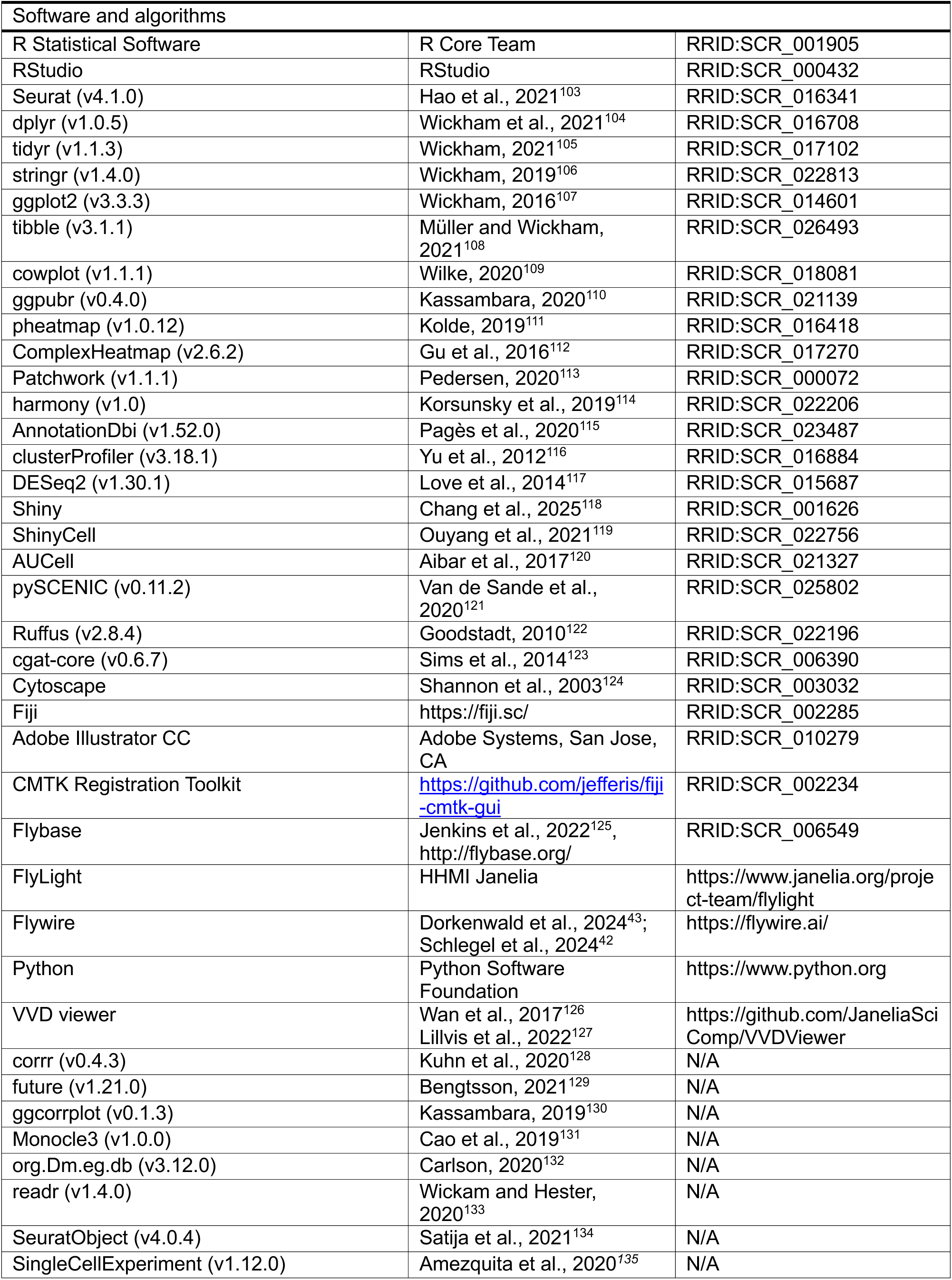
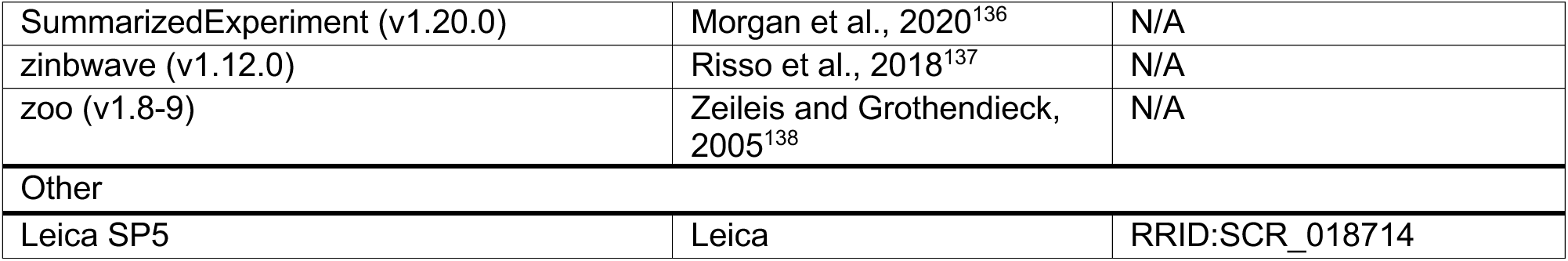

### RESOURCE AVAILABILITY

#### Lead contact

Further information and requests for resources and reagents should be directed to and will be fulfilled by the lead contacts Aaron Allen and Stephen F. Goodwin.

#### Materials availability

All unique/stable reagents generated in this study are available from the lead contacts without restriction.

### EXPERIMENTAL MODEL AND SUBJECT DETAILS

#### Drosophila stocks

All *Drosophila melanogaster* stocks were reared at 25°C and 40-50% humidity on standard cornmeal-agar food with a 12:12 light/dark cycle. Genotypes of the flies used were reported in the figure and legend. All strains used in the study are indicated in the key resources table.

#### Full Genotype List

Full Genotype List:
| Figure | Full genotype |
| --- | --- |
| **2H, S3C** | *yw\*/w\**; 10xUAS-IVS-myr::GFP/ *inv*<sup>p65AD</sup>; *dsx*<sup>DBD</sup>/+<br>*w\*/unc-4*<sup>AD</sup>; 10xUAS-IVS-myr::GFP/+; *dsx*<sup>DBD</sup>/DC335<br>*yw\*/w\**; 10xUAS-IVS-myr::GFP/ *bsh*<sup>p65AD</sup>; *dsx*<sup>DBD</sup>/+<br>*yw\*/w\**; 10xUAS-IVS-myr::GFP/+; *dsx*<sup>DBD</sup>/Fer2<sup>p65AD</sup><br>*yw\*/w\**; 10xUAS-IVS-myr::GFP/+; *dsx*<sup>DBD</sup>/+; *sv*<sup>p65AD</sup>/+<br>*yw\*/w\**; 10xUAS-IVS-myr::GFP/+; *dsx*<sup>DBD</sup>/TfAP-2<sup>p65AD</sup> |
| **2N** | *UAS-mCD8::GFP*/+; *dsx*[DBD], *UAS-mCD8::GFP*/+ |
| **3F, S7E** | *yw\*/w\**; *VGlut*<sup>DBD</sup>/+; 10xUAS-IVS-mCD8::GFP/Ptx1<sup>p65AD</sup> |
| **3K, S7F** | *yw<sup>\*</sup>/w<sup>\*</sup>; dve<sup>DBD</sup>/UAS>mCD8::GFP>Chrimson::tdTomato; fru<sup>FLP</sup>/Ptx1<sup>p65AD</sup>* |
| **4E,G, S8A** | *yw<sup>\*</sup>/w<sup>\*</sup>; 10xUAS-IVS-mCD8::GFP/+; Gad1<sup>DBD</sup>/TfAP-2<sup>p65AD</sup>/+* |
| **4F, S8A,B** | *w<sup>\*</sup>/w<sup>\*</sup>; UAS>stop>myrGFP/+; Gad1<sup>DBD</sup>, fru<sup>FLP</sup>/TfAP-2<sup>p65AD</sup>* |
| **4I, S8C** | *w<sup>\*</sup>/w<sup>\*</sup>; 13XlexAop-IVS- GFP-p10/+; fru<sup>P1.LexA</sup>/fru<sup>F</sup>*<br>*w<sup>\*</sup>/w<sup>\*</sup>; 13XlexAop-IVS- GFP-p10/+; fru<sup>P1.LexA</sup>/fru<sup>M</sup>* |
| **4J, S8E** | *UAS-p35,w<sup>\*</sup>/w<sup>\*</sup>; 10xUAS-IVS-mCD8::GFP/+; Gad1<sup>DBD</sup>/TfAP-2<sup>p65AD</sup>/+* |
| **6D, S12A** | *yw<sup>\*</sup>/w<sup>\*</sup>; UAS>stop>myrGFP/+; dsx<sup>DBD</sup>, Fer2<sup>p65AD</sup>/fru<sup>FLP</sup>/*<br>*yw<sup>\*</sup>/w<sup>\*</sup>; 10xUAS-IVS-myr::GFP/+; dsx<sup>DBD</sup>/CG42458<sup>p65AD</sup>*<br>*yw<sup>\*</sup>/w<sup>\*</sup>; 10xUAS-IVS-myr::GFP/+; dsx<sup>DBD</sup>/Dbx<sup>p65AD</sup>*<br>*yw<sup>\*</sup>/w<sup>\*</sup>; 10xUAS-IVS-myr::GFP/+; dsx<sup>DBD</sup>/CG15537<sup>p65AD</sup>* |
| **S4H** | *Y/w<sup>\*</sup>; UAS-WM/Vmat<sup>p65AD</sup>; fru<sup>DBD</sup>/+* |
| **S6A** | *Y/w<sup>\*</sup>; UAS-WM/+; fru<sup>DBD</sup>/+; sv<sup>p65AD</sup>/+* |
| **S6B** | *Y/w<sup>\*</sup>; 10xUAS-IVS-myr::GFP/+; dsx<sup>DBD</sup>/+; sv<sup>p65AD</sup>/+* |
| **S6D** | *Y/unc-4<sup>AD</sup>; UAS-WM/+; fru<sup>DBD</sup>/DC335* |
| **S6F** | *Y/w<sup>\*</sup>; UAS-WM/+; fru<sup>DBD</sup>/TfAP-2<sup>p65AD</sup>* |
| **S7D** | *w<sup>\*</sup>/w<sup>\*</sup>; VGlut<sup>DBD</sup>/UAS>stop>myrGFP; fru<sup>FLP</sup>/Ptx1<sup>p65AD</sup>* |
| **S7G** | *Y/w<sup>\*</sup>; UAS-WM/+; fru<sup>DBD</sup>/Fer2<sup>p65AD</sup>*<br>*Y/w<sup>\*</sup>; UAS-WM/tsh<sup>VP16AD</sup>; fru<sup>DBD</sup>/+*<br>*Y/w<sup>\*</sup>; UAS-WM/+; fru<sup>DBD</sup>/TfAP-2<sup>p65AD</sup>*<br>*Y/w<sup>\*</sup>; UAS-WM/+; fru<sup>DBD</sup>/Ptx1<sup>p65AD</sup>*<br>*Y/unc-4<sup>AD</sup>; UAS-WM/+; fru<sup>DBD</sup>/DC335*<br>*Y/w<sup>\*</sup>; UAS-WM/+; fru<sup>DBD</sup>/Dbx<sup>p65AD</sup>*<br>*Y/w<sup>\*</sup>; UAS-WM/bsh<sup>p65AD</sup>; fru<sup>DBD</sup>/+*<br>*Y/w<sup>\*</sup>; UAS-WM/inv<sup>p65AD</sup>; fru<sup>DBD</sup>/+*<br>*Y/w<sup>\*</sup>; UAS-WM/+; fru<sup>DBD</sup>/D<sup>VP16AD</sup>*<br>*Y/w<sup>\*</sup>; UAS-WM/+; fru<sup>DBD</sup>/+; sv<sup>p65AD</sup>/+*<br>*Y/w<sup>\*</sup>; UAS-WM/vg<sup>p65AD</sup>; fru<sup>DBD</sup>/+*<br>*Y/w<sup>\*</sup>; UAS-WM/dve<sup>p65AD</sup>; fru<sup>DBD</sup>/+*<br>*Y/w<sup>\*</sup>; UAS-WM/twit<sup>p65AD</sup>; fru<sup>DBD</sup>/+* |
| **S8D** | *w<sup>\*</sup>/w<sup>\*</sup>; 13XlexAop-IVS-GFP-p10/+; fru<sup>P1.LexA</sup>/+* |
| **S8F** | *yw<sup>\*</sup>/w<sup>\*</sup>; 10xUAS-IVS-myr::GFP/ Dh31-R<sup>p65AD</sup>; dsx<sup>DBD</sup>/+*<br>*Y/w<sup>\*</sup>; UAS-WM/ Dh31-R<sup>p65AD</sup>; fru<sup>DBD</sup>/+* |
| **S10A,B** | *Y/w<sup>\*</sup>; UAS-WM/+; fru<sup>DBD</sup>/+; sv<sup>p65AD</sup>/+*<br>*yw<sup>\*</sup>/w<sup>\*</sup>; 10xUAS-IVS-myr::GFP/+; dsx<sup>DBD</sup>/+; sv<sup>p65AD</sup>/+* |

Sexes of all genotypes are stated in the figures.

### METHOD DETAILS

#### Generation of meta-central brain neuronal atlas

The generation of this atlas is described in detail in a companion paper^19^. All code and specific details of what was run, please refer to https://github.com/aaron-allen/Dmel-adult-central-brain-atlas. Briefly, we combined our newly generated scRNA-seq data of the adult central brain^19^ with multiple re-processed publicly available 10x chromium 3’ chemistries datasets^91,93–99,139,140^. These datasets were integrated, and non-neuronal cells, peripheral neurons, and optic lobe neurons were annotated and removed. The remaining central brain neurons were re-clustered, resulting in a well-integrated transcriptional central brain neuron meta-atlas with 329,466 cells.

#### Annotation of sex

Two of the datasets used in our meta-analysis did not separate the sexes into separate samples^93,95^. As a result, we’ve had to rely on in-silico sexing of these samples. We used Seurat’s “AddModuleScore” function with the male-specific genes *lncRNA:roX1* and *lncRNA:roX2*. Cells with a module score less than zero were labelled female, and cells with module scores greater than zero were labelled male. To test the efficacy of this method, we performed *in silico* sexing of the datasets where the sexes were processed separately and achieved a precision of 0.93-1.00 and a recall of 0.89-0.97 for identifying a cell as female, depending on the dataset.

#### Differential gene expression analysis

Differential gene expression between the sexes was conducted in a similar fashion to^95^. We use the zinbwave package (Zero-Inflated Negative Binomial-based WAnted Variation Extraction) to account for dropout and the count-based nature of the data^137^. We included “dataset” as a covariate in the model to correct for differences between the individual datasets. Differential expression was conducted with DESeq2^117^. Even with correcting for dataset in the models for the differential expression testing, multiple genes initially recovered to be sex-biased exhibited strong dataset-specific expression (Figure S2). To eliminate these genes, we filtered the genes by the following criteria: (1) the total UMI of the gene across all cells greater than or equal to 400, (2) the maximum UMI observed within a cell greater than or equal to 4, (3) the percent contribution of UMI from each dataset less than or equal to 60%, and (4) the percent contribution of UMI from the Park et al. (2024) dataset less than or equal to 30%.

#### Gene ontology analysis

Gene ontology analysis was performed using the packages “clusterProfiler“^116^ with “org.Dm.eg.db“^132^ and “ AnnotationDbi“^115^. The “enrichGO” function was used to determine the enriched terms for each of the molecular function, biological process, and cellular compartment categories (Figure S1E).

#### Subclustering doublesex and fruitless neurons

To subcluster *dsx+* neurons, all neurons with greater than or equal to 1 UMI of *dsx* expression were extracted. For our generated dataset^19^, we transgenically expressed *EGFP* using *dsx^Gal^*^4^, and so we also included any cell with at least 1 UMI of *EGFP* from our dataset. For *fru+* subcluster, dataset specific thresholds were chosen (Davie-2018 > 1; Baker-2021 > 2; Park-2022 > 2; Dopp-2024 > 3; LeeBenton-2023 > 3; Li-2022 > 3; Lu-2023 > 3; Park-2024 > 3). These cells were extracted and re-integrated using the “harmony” package^114^ using “RunHarmony”, correcting for dataset of origin, preparation type (cell vs. nuclei), and individual sample ID metadata. The re-integrated cells were clustered with “FindNeighbors” and “FindClusters” using the Louvain algorithm.

#### Early-vs late-born morphology quantification

To annotate early-born (“Imp>dati”) and late-born (“dati>Imp”), we used Seurat’s “AddModuleScore” to calculate the average expression level of early (*Imp*, *mamo*) and late (*dati*, *pros*) gene programs across all cells. A cell was annotated as early-born if the l2-normalised early-born module score was both greater than zero and greater than the l2-normalised late-born module score. Conversely, a cell was annotated as late-born if the l2-normalised late-born module score was both greater than zero and greater than the l2-normalised early-born module score.

#### Gene regulatory network analysis with SCENIC

Gene regulatory network analysis was performed as previously described^19^. Briefly, pySCENIC^121^ (v0.11.2) was run on select, individual subclustered clusters to identify transcription factor based regulons by using the gene expression data as input. The raw expression matrix was filtered, modules comprising transcription factors and co-expressed genes were generated using GRNBoost2, and then pruned to remove indirect targets lacking transcription factor motif using cisTarget. Cells were then scored for the activity of each high-confidence regulon using the AUCell^120^. Cell-type enriched regulons were calculated using Seurat’s “FindAllMarkers” function. Gene regulatory networks were visualized with Cytoscape (Shannon et al., 2003). For visualization purposes, GRNs were further filtered to only include positive regulons less than 100 genes in size, and each node (gene) had to have a maximum expression of at least 3 UMI and total expression of at least 80 UMI.

#### Pseudotime analysis

Pseudotime analysis was performed using Monocle3^131^ (v1.0.0) using standard methods. Briefly, the data were processed with functions “new_cell_data_set”, “preprocess_cds”, “align_cds”, “reduce_dimension”, “cluster_cells”, and “learn_graph” using default settings. Cells were ordered with “order_cells” setting all *Imp*+ cells as the root. Genes with significantly variable expression across pseudotime were calculated with “graph_test” and filtered for q-value less than 1e-50 and Moran’s I greater than 0.2. Heatmaps of significantly varying genes were plotted with the package “ComplexHeatmap“^112^.

#### Re-clustering and correcting for early-vs late-born neurons

In our companion manuscript^19^, we show that early-born *Imp*+ neurons have separated “punctate” like clustering as compared to the late-born *dati+* neurons. We also showed that these late-born unsupervised clusters may correspond to hemilineages. To better approximate hemilineage and include the appropriate early-born and late-born neurons with unsupervised clusters, we projected early-vs late-born into a shared embedding using the “harmony” package. This re-clustered embedding was used to approximate hemilineage identity (Figure 3I; Figure 3J; Figure 4K; Figure 5F).

#### Generating a web app

Our interactive web tool for visualization of these scRNA-seq data was generated using a modified version of “ShinyCell”, an R-based Shiny App^119^; https://github.com/SGDDNB/ShinyCell). Specifically, we forked and modified “easyshiny” (https://github.com/NBISweden/easyshiny), which is a forked version of the original “ShinyCell”.

#### Generation of split-Gal4 driver lines

We used existing coding intronic Minos-mediated integration cassette/CRISPR-mediated integration cassette (MiMIC/CRIMIC) lines^141,142^ (see Key Resources Table) to generate split-GAL4 drivers using the Trojan method^143^. pBS-KS-attB2-SA()-T2A-Gal4DBD-Hsp70 or pBS-KS-attB2-SA()-T2A-p65AD-Hsp70 vectors^143^ with the appropriate reading frame were inserted in the MIMIC/CRIMIC locus of a given line via recombinase-mediated cassette exchange through injection (BestGene, CA). Stocks were generated with the transformed flies as described in^143^.

#### Immunohistochemistry

After a brief pre-wash of adult flies in 100% EtOH to remove hydrophobic cuticular chemical compounds, brains were dissected in PBS at RT (20-25°C), collected in 2 mL sample tubes and fixed with 4% formaldehyde (Sigma-Aldrich) in PBS (Sigma-Aldrich) for 20 min at RT. After fixation, tissues were washed in 0.5-0.7%PBS/Triton X-100 (Sigma-Aldrich) (PBT) 3 times each for 20 min at RT. After blocking in 10% normal goat serum (Sigma-Aldrich) in PBT (NGS/PBT) overnight (8-12 h) at RT, tissues were incubated in primary antibody solutions for 48-72 hrs at 4°C (1:1000, rabbit anti-GFP, Thermo Fisher Scientific; 1:1000, chicken anti-GFP, Abcam; 1:1000, rabbit anti-RFP, Antibodies Online; 1:10, mouse anti-Brp, Developmental Studies Hybridoma Bank; 1:500, rabbit anti-Dh31, Lin et al., 2022; 1:500, guinea pig anti-Otp, Hildebrandt et al., 2020). After 4 washes in PBT for 1 h each at RT, tissues were incubated in secondary antibody solutions for 48 h at 4°C with the exception of anti-Dh31 staining, in which tissues were incubated in secondary antibody solution for 24 h at 4°C (1:500, anti-rabbit Alexa Fluor 488, anti-rabbit Alexa Fluor 633, anti-chicken Alexa Fluor 488, anti-mouse Alexa Fluor 546, anti-rat Alexa Fluor 633, anti-guinea pig Alexa Fluor 633, Thermo Fisher Scientific). After 4 washes in PBT for 1 h each at RT specimens were imaged directly or 70% glycerol in PBS was added to the sample tubes, which were subsequently transferred to -20°C and kept for at least 8 hr for tissue clearing. Specimens were mounted in Vectashield (Vector Laboratories).

#### Confocal image acquisition and processing

Confocal image stacks were acquired on a Leica TCS SP5 confocal microscope at 1024 × 1024-pixel resolution with a slice size of 0.29 µm or 1 µm. Water-immersion 25x and oil-immersion 40x objective lenses were used for brain images. Images were registered onto the unisex template brain using the Fiji Computational Morphometry Toolkit (CMTK) Registration GUI (https://github.com/jefferis/fiji-cmtk-gui). For segmented images, we used the software VVD Viewer^126,127^ (https://github.com/JaneliaSciComp/VVDViewer) to render the registered image stacks in 3D, manually mask other neurons co-labelled in the image, and segment out neurons of interest. All segmented images include unsegmented versions within the supplemental figures.

## DATA and CODE AVAILABILTIY

The code used in this analysis will be made available upon publication. The following data set was generated in our companion manuscript: (to follow upon publication). The following previously published data sets were used: GSE107451, GSE152495, GSE207799, GSE221239, GSE247965, E-MTAB-10519, GSE218661, GSE261656, GSE142789, GSE153723, GSE218257, GSE162098, and GSE160370. Any additional information needed to reanalyse the data reported in this paper is available from the lead contact by request. The datasets generated and described here can be visualized and explored at https://www.flycns.com/. Any additional information needed to reanalyse the data reported in this paper is available from the lead contact by request. The following previously published data sets were used:

## Acknowledgments

We thank E. Rideout, E. J. Clowney, Y. Ding, J. Walsh, and the Goodwin Lab for helpful discussions and critical reading of the manuscript. We are grateful to F. Casares, C. Desplan, T. Erclik, J. Hall, H. Lacin, G. Miesenböck, M. Nitabach, T. Shirangi, S. Waddell, U. Walldorf, and J. Wang for sharing important stocks and reagents with us. Stocks obtained from the Bloomington Drosophila Stock Center (NIH P40OD018537) were used in this study. A.M.A. and M.C.N. were supported by a Wellcome Trust Senior Investigator Award to S.F.G (106189/Z/14/Z) and a grant from the BBSRC (BB/X016595/1) to M.C.N, A.M.A., and S.F.G. T.N. and F.A. were supported by a grant from the BBSRC (BB/T001348/1) to T.N. and S.F.G. A.M.A was supported by a Wellcome Collaborative Award to S.F.G. (209235/Z/17/Z).

## Author contributions

Conceptualization, A.M.A., M.C.N., and S.F.G.; Methodology, A.M.A., M.C.N., T.N., F.A., and S.F.G.; Investigation, A.M.A., M.C.N., T.N., and F.A.; Resources, S.F.G.; Writing, A.M.A., M.C.N., T.N., F.A., and S.F.G.; Supervision, S.F.G.; Funding Acquisition, S.F.G.

